# CARM1-mediated methylation controls interactions of ALIX with key partners important for cytokinesis

**DOI:** 10.1101/2025.07.06.663356

**Authors:** Solène Huard, Samyuktha Suresh, Jessica Vayr, Sonia Ruggiero, Rayan Dakroub, Vanessa Masson, Anđela Petrović, Rayane Dibsy, Julie Thomsen, Yara Hajj Younes, Clémence Vissotsky, Louise Amara, Lucid Belmudes, Christine Chatellard, Rémy Sadoul, Ahmed El Marjou, Yohann Couté, Damarys Loew, Raphaël Guérois, Nicolas Reynoird, Arnaud Echard, Thierry Dubois

## Abstract

CARM1 is an arginine methyltransferase with a well-established role in regulating gene expression, but its cytoplasmic functions remain largely uncharacterized. Here, we identified ALIX, a protein acting with the ESCRT-III machinery in numerous membrane remodeling events, as a main cytoplasmic partner and a relevant substrate of CARM1. We demonstrate that CARM1 methylates arginine residues within the proline-rich motif of ALIX. At the molecular level, ALIX methylation impairs SH3-dependent association with CD2AP, CIN85, and endophilin-A2 that are required for cytokinesis. Using a mutant of ALIX that is unable to bind these proteins, we further show that these interactions are essential for ALIX functions during cytokinesis. Altogether, this work highlights the significance of arginine methylation as a regulatory post-translational modification important for the final step of cell division, by modulating interactions between proline-rich domains and their SH3-containing partners.

## Introduction

Arginine methylation is a post-translational modification (PTM) catalyzed by the nine members of the protein arginine methyltransferase (PRMT) family^1-4^. The prevailing view is that protein methylation triggers interaction with specific partners known as “readers”, which propagate downstream signaling events^5^. By methylating nuclear proteins including histones and transcription factors, PRMTs are involved in various cellular processes such as the regulation of transcription^1-4^. PRMT4, also known as coactivator-associated arginine methyltransferase 1 (CARM1), catalyzes both monomethylation (MMA, me1) and asymmetric dimethylation (ADMA, me2a) of arginine residues^6-8^. CARM1 is distinguished from other PRMTs by its specificity for methylating arginine residues within proline-rich motifs^9-12^. Elevated levels of CARM1 have been observed in various cancer types, suggesting its potential as a therapeutic target and a diagnostic marker^6-8^. CARM1 is found in the nucleus and is mostly studied in the context of gene regulation^13^, influencing processes such as development and autophagy^6-8^. Importantly, CARM1 is also localized in the cytoplasm^6-8^, although its role in that cellular compartment remains elusive.

Cytokinesis is the last step of cell division, leading to the separation of two daughter cells connected by an intercellular bridge filled with microtubules^14-16^. At the center of this bridge lies the midbody, where proteins required for the completion of cytokinesis are recruited. The final abscission of the intercellular bridge requires local F-actin remodeling and microtubule severing as well as the endosomal sorting complex required for transport (ESCRT) machinery^14-22^. ESCRT-I proteins (TSG101 and VPS37) and ESCRT-III proteins (CHMP4B) directly associate with ALIX (ALG-2 interacting protein X), also known as programmed cell death 6 interacting protein (PDCD6IP). ALIX plays an important role in the late steps of cytokinesis by contributing to the recruitment of ESCRT-III to the midbody^23-27^. It acts in parallel to TSG101, and both ALIX and TSG101 are recruited to the midbody by CEP55^24,25,27,28^. In addition, ALIX bridges and stabilizes the ESCRT-III filaments with the plasma membrane by interacting with the syntenin/syndecan-4 complex, ensuring proper stability of ESCRT-III at the abscission site^26^. ALIX consists of several domains, including an N-terminal Bro1 domain that mediates interaction with capping proteins and CHMP4B, a central V domain that binds to the syntenin/syndecan-4 complex, and a C-terminal proline-rich domain (PRD) responsible for binding to CEP55, CD2AP (and its homologue CIN85), endophilins, TSG101, Src and ALG2^23,25,29-35^ (Supplementary Fig. 1). Among these ALIX partners, many have been involved in cytokinesis, notably CEP55, TSG101, CD2AP, CIN85, capping proteins and endophilin A2^15,23-27,36-40^. Human ALIX has been shown to be phosphorylated, ubiquitinated and palmitoylated. The phosphorylation of ALIX leads to a change of its conformation into an open and activated form which is required for proper cytokinesis^41^. Whether ALIX is regulated by other PTMs is currently unknown.

Here, we identified ALIX as a major partner of CARM1 and found that the methyltransferase domain of CARM1 interacts with the PRD of ALIX. We showed that CARM1 methylates the PRD of ALIX at three arginine residues (R_745_, R_757_, R_767_) *in vitro* and at least at two of them (R_745_, R_757_) in cells. Searching for their methyl-sensitive protein interactors, we found that CD2AP, CIN85, and endophilin-A2, previously reported to associate to ALIX through their SH3 domains^23,25,29,42^, indeed bound to ALIX but only when these arginine residues were not methylated, highlighting the critical role of arginine methylation in preventing association with their SH3 domains. Importantly, a mutant form of ALIX that is unable to interact with these proteins failed to rescue cytokinetic defects caused by the loss of endogenous ALIX. Altogether, we propose that CARM1-mediated methylation of ALIX regulates interactions with key partners that are crucial for its function in the completion of cytokinesis. Furthermore, our study identifies arginine methylation as a previously unrecognized post-translational modification important for cytokinesis.

## Results

### The cytoplasmic protein ALIX is a main partner of CARM1

To get insight about CARM1 cytoplasmic functions, we characterized its interactome by immunoprecipitating endogenous CARM1 from several breast cancer cell lines and HeLa cells, followed by mass spectrometry (MS)-based proteomic analysis. Remarkably, ALIX was consistently retrieved among the top five CARM1 partners in all cell lines, using three different CARM1 antibodies (Fig. 1a, Supplementary Fig. 2), even under high stringency conditions (1% NP-40, Supplementary Fig. 2). Two isoforms of CARM1 are expressed in mammalian cells, CARM1-FL (Full-Length) and CARM1-Δ15, the latter being the predominant isoform in most tissues and cell lines^7,8,43-45^. Immunoprecipitation of ectopically expressed Flag-CARM1-FL or Flag-CARM1-Δ15 from HEK-293T cells, followed by MS-based proteomic analyses, identified a very limited number of potential partners, with ALIX emerging as the top candidate for both isoforms (Fig. 1b). Conversely, immunoprecipitation of endogenous ALIX in breast cancer cells and HeLa cells followed by MS-based proteomic analysis retrieved CARM1 in addition to well-established ALIX partners such as CD2AP, CIN85, endophilin-A2, CEP55 and CHMP4B (Fig. 1c). Endogenous CARM1/ALIX interaction was confirmed by immunoprecipitating CARM1 followed by western blot analysis (Fig. 1d). The association was further validated by co-expressing Flag-CARM1 and GFP-ALIX in HEK-293T cells (Fig. 1e). Moreover, ALIX interacted specifically with CARM1, and not with PRMT1, PRMT2, PRMT3 and PRMT5 (Fig. 1e). Altogether, these results identify ALIX as a main partner of both CARM1 isoforms. Furthermore, ALIX specifically interacts with CARM1 among the PRMT family members analyzed.

**Figure 1.**
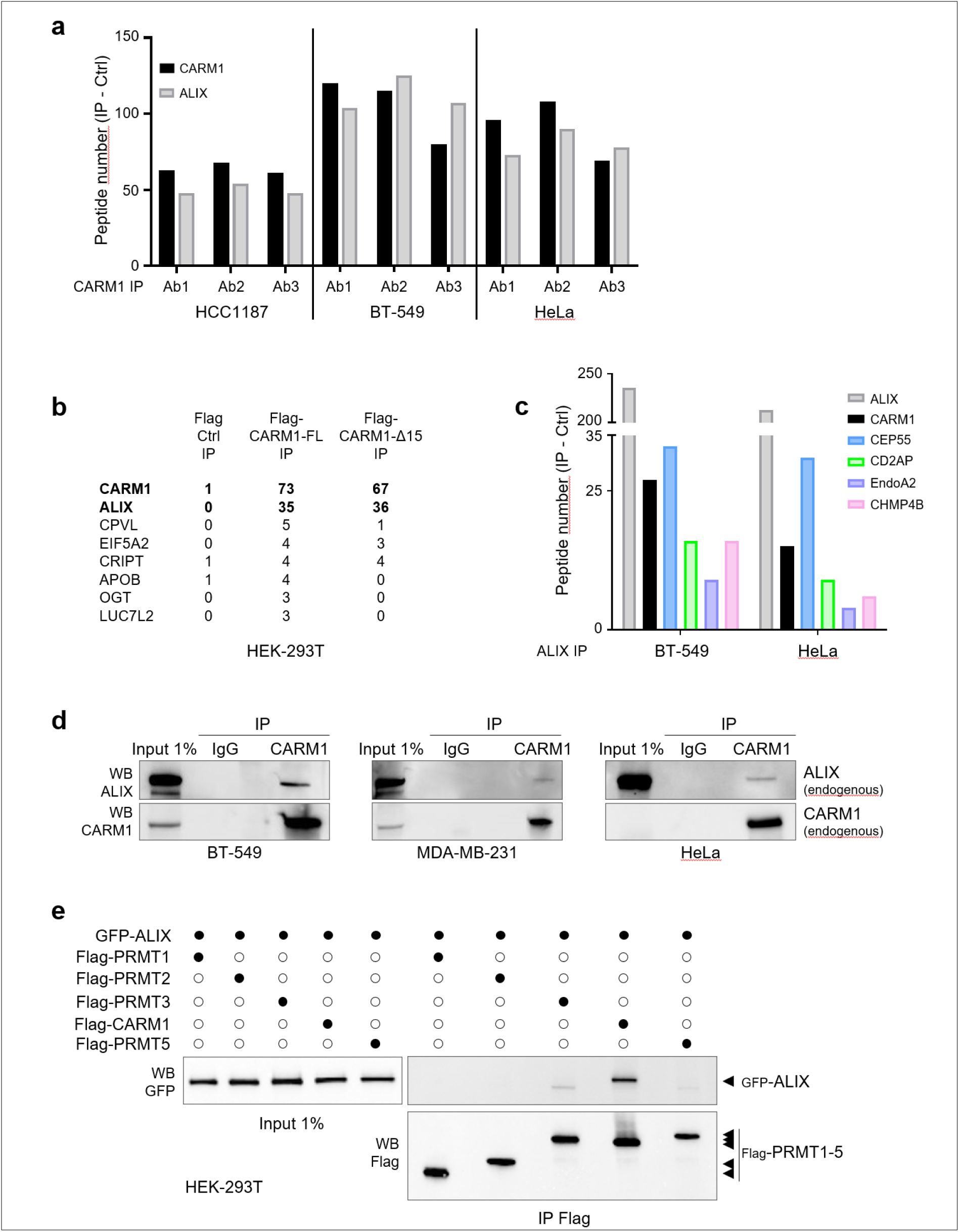
ALIX is a main cytoplasmic partner of CARM1. **(a)** Peptide counts (CARM1 IP – IgG IP) detected by MS-based proteomic analysis for CARM1 and ALIX following CARM1 immunoprecipitation (IP) using three different CARM1 antibodies (Ab1-Ab3) from breast cancer cells (HCC1187, BT-549) and HeLa cells. The cytoplasmic protein ALIX was retrieved as one of the top CARM1 partners in all cell lines. **(b)** Peptide counts corresponding to proteins identified by MS-based proteomic analysis following IP using a Flag antibody of ectopically expressed Flag-CARM1-FL and Flag-CARM1-Δ15 in HEK-293T cells. The complete list of retrieved proteins is shown (CARM1-FL IP – IgG IP > 3). **(c)** Peptide counts (ALIX IP – IgG IP) detected by MS-based proteomic analysis for ALIX and known interactors as well as for CARM1 following ALIX IP in BT-549 and HeLa cell lines. **(d)** Endogenous CARM1 was immunoprecipitated from BT-549, MDA-MB-231 or HeLa cells and the interaction with ALIX was assessed by western blot (*N* = 3). **(e)** HEK-293T cells were co-transfected with plasmids as indicated. IP using a Flag antibody was performed and the interaction between GFP-ALIX and Flag-PRMTs were assessed by western blot. Images shown are representative of three independent experiments.

### The catalytic domain of CARM1 interacts with the proline-rich domain of ALIX

ALIX is composed of an N-terminal Bro1 domain, a central V-domain, and a C-terminal intrinsically disordered proline-rich domain (PRD), all serving as platforms for protein-protein interactions (Fig. 2a, Supplementary Fig. 1). To assess which domain(s) of ALIX interact with CARM1, we designed mCherry-tagged ALIX constructs expressing ALIX full-length (FL), ALIX-Bro1, ALIX-ΔBro1, ALIX-V, or ALIX-PRD (Fig. 2a) and co-transfected them with Flag-CARM1 (which refers from now to CARM1-FL) in HEK-293T cells, followed by Flag immunoprecipitation. As expected, ALIX-FL co-immunoprecipitated with Flag-CARM1 (Fig. 2b). ALIX-Bro1 did not bind to CARM1, while ALIX-ΔBro1 interacted with CARM1, indicating that the Bro1 domain is not important for the association with CARM1 (Fig. 2b). ALIX-PRD but not ALIX-V bound to CARM1 (Fig. 2c), showing that CARM1 interacts with the PRD of ALIX. Confirming this finding, a mutant of ALIX lacking the PRD (ALIX-ΔPRD) did not bind to endogenous CARM1 (Fig. 2d, Supplementary Fig. 3). These results demonstrate that the PRD domain is necessary and sufficient for CARM1 binding to ALIX.

**Figure 2.**
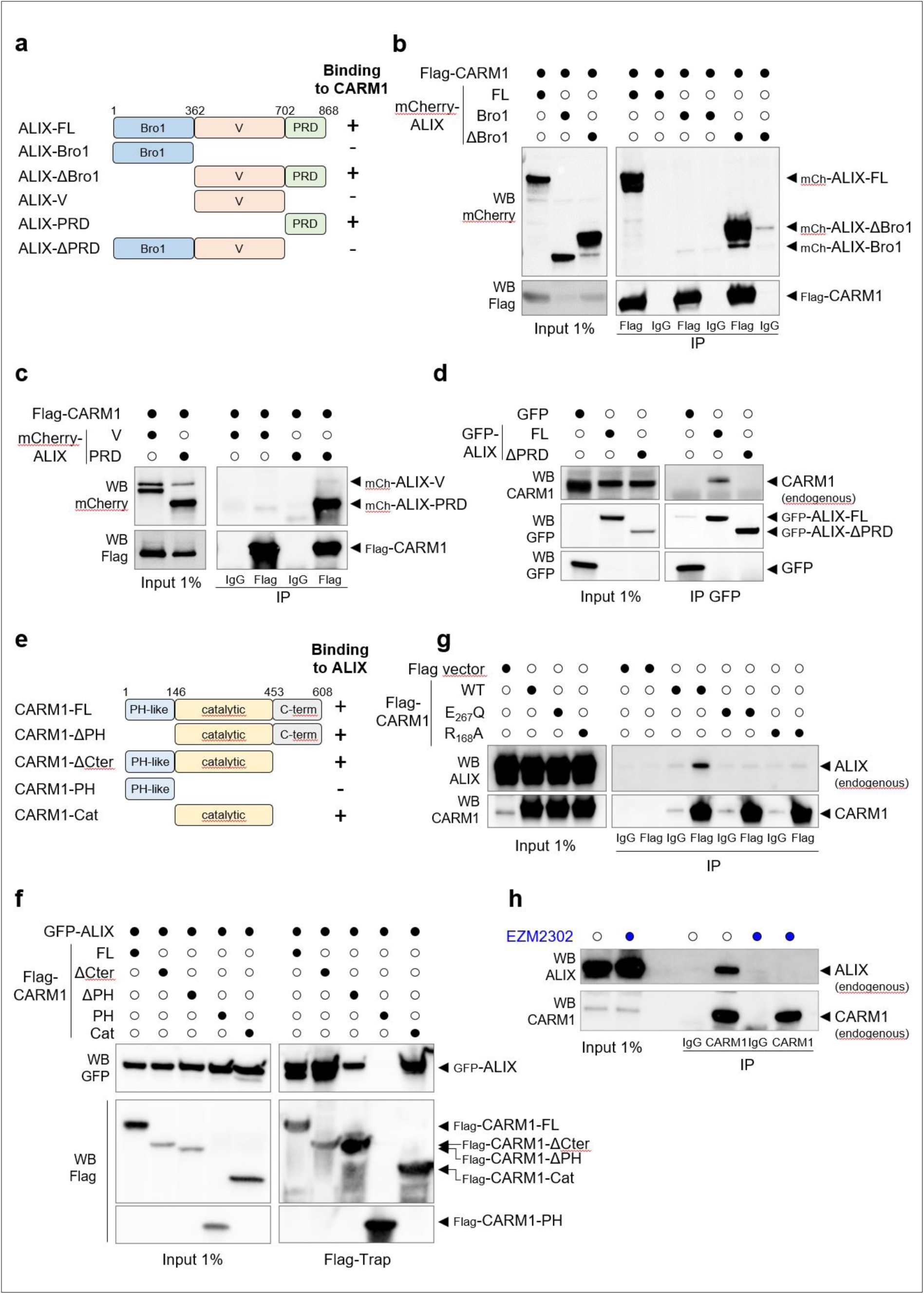
The catalytic domain of CARM1 associates with the proline-rich domain of ALIX. **(a)** Schematic representation of the different domains of ALIX, the different mutants used in B-D, and their ability to bind to CARM1. (**b,c)** HEK-293T cells were co-transfected with the constructs as indicated. Immunoprecipitation (IP) using a Flag antibody was performed and the interaction between Flag-CARM1 and the different mCherry-ALIX proteins were assessed by western blot *(N = 3)*. **(d)** HEK-293T cells were transfected with the constructs as indicated. IP using a GFP antibody was performed and the interaction between endogenous CARM1 and the different GFP-ALIX proteins were assessed by western blot (*N = 4*). **(e)** Schematic representation of the different domains of CARM1, the different mutants used in Fig. 2f, and their ability to bind to ALIX. (**f)** HEK-293T cells were co-transfected with the constructs as indicated. IP using Flag-Trap was performed and the interaction between GFP-ALIX and the different Flag-CARM1 proteins were assessed by western blot *(N = 3)*. **(g)** BT-549 cells were transfected with the indicated constructs. IP using an anti-Flag antibody was performed and the interaction between endogenous ALIX and Flag-CARM1 was assessed by western blot *(N = 3)*. **(h)** BT-549 cells were incubated with DMSO or 5 µM EZM2302 for 48 h. IP of endogenous CARM1 was performed and its interaction with endogenous ALIX assessed by western blot (h) *(N = 3)*. The images are representative of at least three independent experiments (b-d, f-h).

CARM1 is composed of an N-terminal PH-like domain, a central catalytic domain, and a C-terminal domain (Fig. 2e). To investigate which CARM1 domains are interacting with ALIX, we generated truncated mutants of Flag-tagged CARM1 and expressed them in HEK-293T cells (Fig. 2f). Deleting either the PH-like domain or the C-terminus domain of CARM1 did not affect its interaction with GFP-ALIX, suggesting that the catalytic domain of CARM1 is sufficient to associate with ALIX (Fig. 2f). Indeed, the catalytic domain of CARM1 alone bound to GFP-ALIX, while the PH-like domain alone did not (Fig. 2f). Since the catalytic domain of CARM1 is important for ALIX interaction, we wondered whether enzyme-dead mutants of CARM1 were able to interact with ALIX. Mutation of CARM1 in either the methyl-donor SAM binding site (CARM1-R_168_A)^46^ or in the substrate arginine binding site (CARM1-E_267_Q)^47^ did not associate with ALIX (Fig. 2g). In addition, treating cells with a substrate competitive CARM1 inhibitor (EZM2302)^48^ prevented CARM1 interaction with ALIX in cells (Fig. 2h). This suggests that an intact and free catalytic pocket of CARM1 is required for its association with ALIX. Collectively, these biochemical data show that the catalytic domain of CARM1 associates with the PRD of ALIX.

### ALIX is a substrate of CARM1 in cells and *in vitro*

We next investigated whether ALIX could serve as a substrate for CARM1. Large-scale studies characterizing the PRMT methylome have reported that ALIX could be methylated at multiple arginine residues in cells^10,11,49-54^: R_322_ (Bro1 domain), R_456_ and R_606_ (V domain), and R_745_, R_757_ and R_767_ (PRD) (Fig. 3a). The three arginine residues within the PRD of ALIX, which are conserved across species (Fig. 3b), were the most reported in these studies, and R_745_ was the most frequent (Fig. 3a). Since CARM1 is the only PRMT specifically methylating arginine residues within proline rich motifs^9-12^, the aforementioned arginine residues represent potential targets for CARM1-mediated methylation (Fig. 3b).

**Figure 3.**
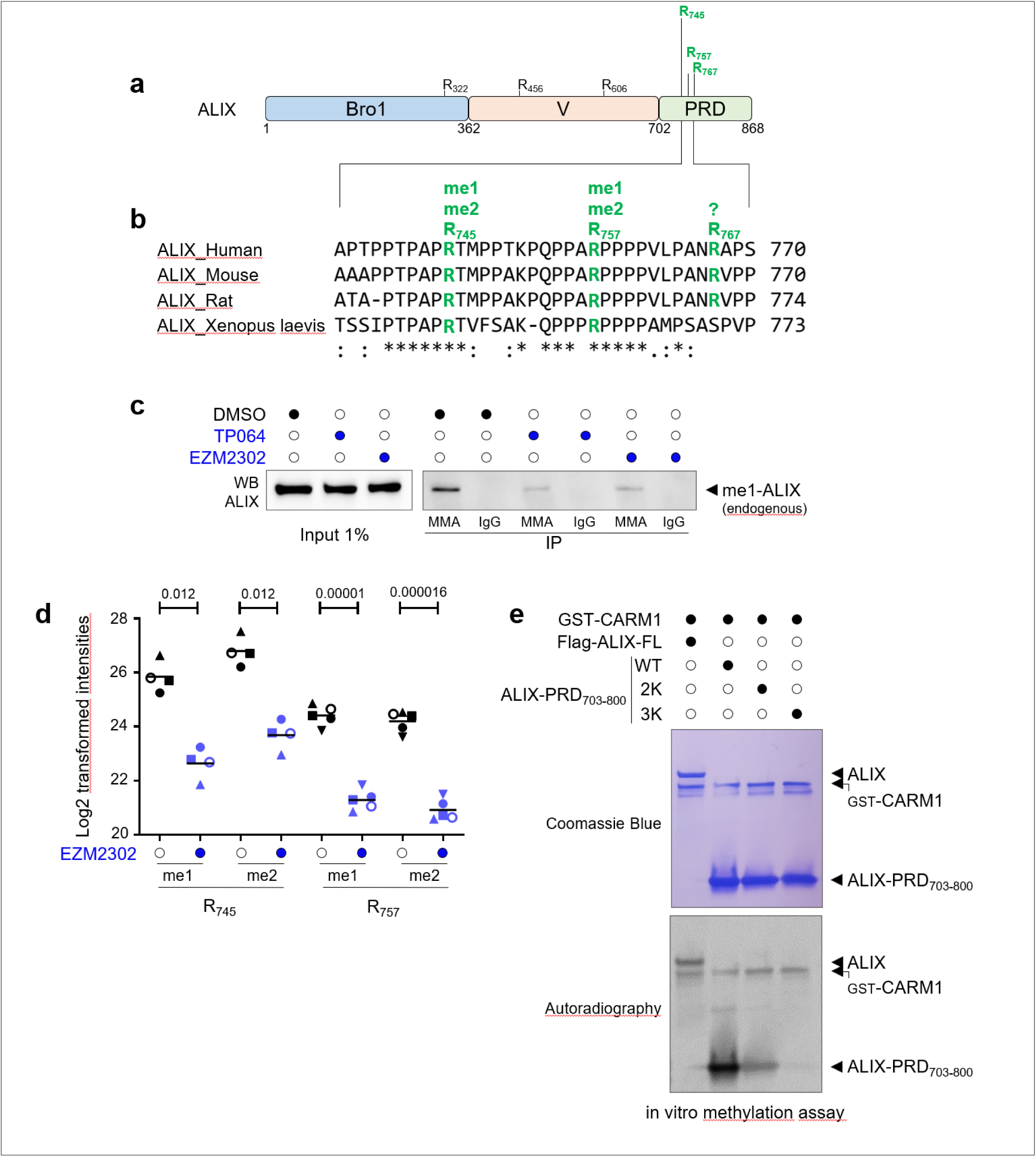
CARM1-dependent methylation of ALIX within its proline-rich domain. **(a)** Schematic representation of the different domains of ALIX and the methylated arginine residues identified by large-scale MS-based proteomic analyses^10,11,49-54^. The most frequent residues that were identified are in the PRD and indicated in green. **(b)** Amino-acid sequence surrounding the three arginine residues of the PRD domain of ALIX found to be methylated (in green) in large-scale MS-based proteomic analyses. These residues and those in proximity are conserved across species. We were not able to detect methylation of R_767_ (indicated by “?”). An asterisk (*) indicates positions that are 100% conserved in the alignment, a colon (:) indicates conservation between groups of strongly similar properties, while a period (.) indicates conservation between groups of weakly similar properties. **(c)** Protein lysates from BT-549 cells treated with 5 µM CARM1 inhibitors (TP064, EZM2302) or with DMSO for 48 h were subjected to immunoprecipitation (IP) using monomethylarginine (MMA) or IgG antibodies. ALIX monomethylation was assessed by western blot. Images are representative of two independent experiments. (**d)** MS-based graphical representation of Log_2_ transformed intensities of monomethylated (me1) and dimethylated (me2) R_745_ and R_757_ peptides after immunoprecipitation of endogenous ALIX from HeLa cells treated with 1 µM EZM2302 or with DMSO (0.01%) for 48 h, and lysarginase digestion (*N* = 4 for R_745_; *N* = 5 for R_757_). *p<0.05; ****p<0.0001 (Student’s t-test). **(e)** Recombinant GST-CARM1 was incubated in the presence of purified Flag-ALIX-FL or recombinant ALIX-PRD_703-800_ (WT, 2K or 3K mutants) in the presence of SAM [^3^H]. Samples were then subjected to SDS-PAGE. Upper panel, Coomassie stain of proteins in the reaction. Bottom panel, autoradiogram (*N* = 3). CARM1 was automethylated, as described^43,94^.

We first examined whether endogenous ALIX was methylated in our cellular models. Monomethylated proteins were immunoprecipitated using a pan monomethylarginine (MMA) antibody in BT-549 (Fig. 3c) and HeLa (Supplementary Fig. 4a) cells followed by western blot using an anti-ALIX antibody. Endogenous ALIX was found to be monomethylated (Fig. 3c, Supplementary Fig. 4a). A prior incubation of cells with two specific CARM1 inhibitors, EZM2302^48^ or TP064^55^, reduced ALIX monomethylation signals, suggesting that CARM1 monomethylates ALIX in breast cancer cells (BT-549) and HeLa cells (Fig. 3c, Supplementary Fig. 4a). The reverse experiment, starting with the immunoprecipitation of endogenous ALIX and blotting with a pan MMA antibody, confirmed these findings (Supplementary Fig. 4b). We could not use a similar approach to assess whether ALIX undergoes arginine dimethylation in cells due to the lack of pan-ADMA antibodies specific for CARM1 substrates. Therefore, endogenous ALIX was immunoprecipitated and subjected to MS-based proteomic analysis to identify whether ALIX is dimethylated in cells. We demonstrated that ALIX is monomethylated and dimethylated in cells on two arginine residues, R_745_ and R_757_ (Supplementary Fig. 5 and 6). We were unable to determine whether the third arginine of the PRD (R_767_) was methylated in our cells (Fig. 3b), possibly due to technical limitations of peptide identification in MS-based proteomic experiments. Importantly, treatment of cells with EZM2302 reduced the level of monomethylation and dimethylation on R_745_ and R_757_ of endogenous ALIX (Fig. 3d), indicating that CARM1 is responsible for their methylation in cells. In these experiments, the efficacy of CARM1 inhibition by EZM2302 was verified by western blot prior to performing MS-based proteomic analysis (Supplementary Fig. 4c).

We next investigated whether CARM1 can directly methylate ALIX. Radioactive *in vitro* methylation assays showed that CARM1 methylates both ALIX-FL and ALIX-PRD_703-800_ (Fig. 3e). The level of methylation was reduced for ALIX-PRD_703-800_ when R_745_ and R_757_ arginine residues were mutated to lysine (R_745_K, R_757_K; termed ALIX-PRD_703-800_-2K) (Fig. 3e). Furthermore, CARM1 did not methylate ALIX-PRD_703-800_ when the three arginine residues were substituted to lysine (R_745_K, R_757_K, R_767_K; termed ALIX-PRD_703-800_-3K) (Fig. 3e), indicating that CARM1 methylates these three arginine residues *in vitro*. Altogether, these results demonstrate that CARM1 mediates monomethylation and dimethylation of ALIX in cells, on at least two arginine residues (R_745_, R_757_) located in its PRD. *In vitro*, CARM1 methylates R_745_, R_757_, and R_767_ in the PRD of ALIX.

### CARM1-dependent methylation of the proline-rich domain of ALIX impairs SH3-mediated interactions with CD2AP, CIN85 and endophilin-A2

To identify methyl-sensitive protein interactors of the asymmetric dimethylated arginine residues R_745_, R_757_ and R_767_ in ALIX, we generated unmethylated (me0) and asymmetrically dimethylated (me2a) synthetic peptides centered around these amino acids. A stable isotope labeling by amino acids in cell culture (SILAC)-based quantitative proteomic screen using breast cancer cell (MDA-MB-231) extracts was then performed to identify proteins that differentially bind to unmethylated versus asymmetrically dimethylated peptides (Fig. 4a and b, Supplementary Fig. 7). Interestingly, MS-based quantitative proteomic analysis identified CD2AP and its homologue CIN85 as the top candidates specifically associated with the unmethylated R_745_ peptide (Fig. 4a), and endophilin-A2 as the top candidate binding to the unmethylated R_757_ peptide (Fig. 4b). No candidate with such high ratio clearly emerged to specifically pulldown with unmethylated or asymmetrically dimethylated R_767_ peptides (Supplementary Fig. 7). Capping proteins (CAPZA1, CAPZA2 and CAPZB) and syndecan-4 (SDC4) also emerged as candidates specifically associating with unmethylated R_745_ peptide (Fig. 4A) and unmethylated R_757_ peptide (Fig. 4b), respectively. Since capping proteins interact with CD2AP/CIN85^56-59^, they likely bind to the unmethylated R_745_ peptide through their association with CD2AP/CIN85. Interactions of CD2AP, CIN85, capping proteins and endophilin-A2 with the unmethylated arginine residues of ALIX were confirmed by peptide pulldown assays using MDA-MB-231 and HeLa cell lysates followed by immunoblotting (Fig. 4c).

**Figure 4.**
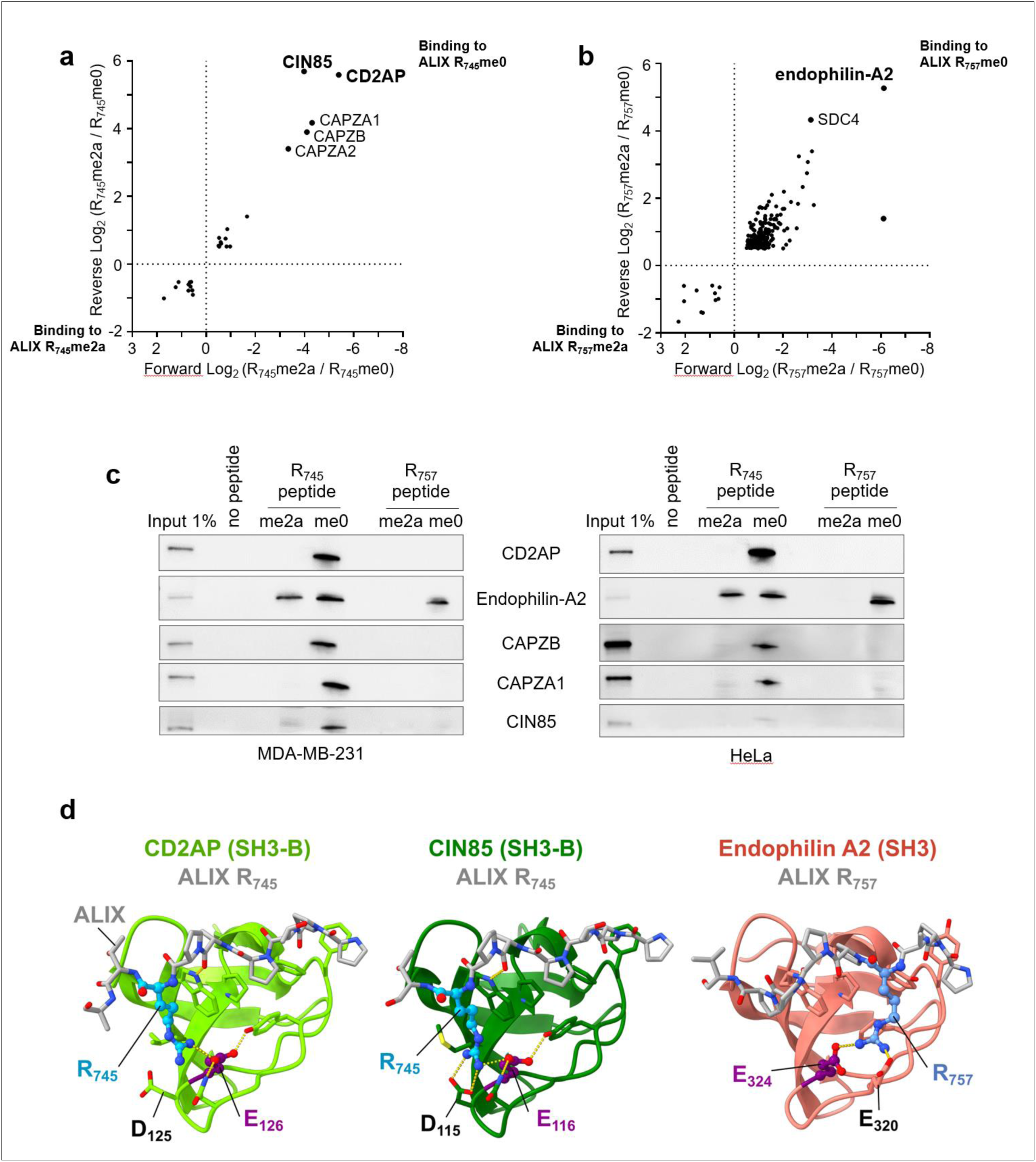
CARM1-dependent methylation of the proline-rich domain of ALIX impairs SH3-mediated interactions with CD2AP, CIN85 and endophilin-A2. **(a,b)** Peptide pull down assays. Proteins interacting with unmethylated (me0) or asymmetrically dimethylated (me2a) R_745_ (a) or R_757_ (b) peptides were identified using MDA-MB-231 cell lysates using SILAC-based quantitative proteomics screen. A scatter plot of the Log_2_ transformed protein-normalized methyl-sensitive peptide SILAC ratios. The x-axis shows the Log_2_ ratio of proteins binding to dimethylated (me2a) versus unmethylated (me0) arginine residues (Forward). The y-axis shows the Log_2_ ratio of a label-swap replicate experiment (Reverse). Specific interactors of unmethylated and asymmetrically dimethylated ALIX reside in the upper right and in the lower left quadrant, respectively. The experiment was done twice with similar results. **(c)** Western blot analysis of protein pulled down using unmethylated or asymmetrically dimethylated ALIX peptides surrounding R_745_ or R_757_ using MDA-MB-231 (left panel) or HeLa cell extracts (right panel) (*N* = 3). Images shown are representative of three independent experiments. **(d)** AlphaFold2 based structural models of the complexes between either R_745_ or R_757_ ALIX peptides and the SH3 domains of CD2AP (left), CIN85 (middle) and endophilin-A2 (right) reveal a key salt-bridge between the side chains of R_745_ or R_757_ and a conserved glutamic acid (E, shown in pink), with another nearby acidic residue (D or E, shown in bold black) likely to contribute favorably to the interaction with the SH3 domains. Details on peptide lengths, SH3 domains boundaries, and model confidence scores are provided in Supplementary Fig. 9.

Remarkably, this unbiased peptide pull-down approach identified CD2AP/CIN85 and endophilin-A2, previously reported to associate through their Src Homology-3 (SH3) domains, with sequences of ALIX containing R_745_ and R_757_ residues, respectively^23,25,29,42^ (Supplementary Fig. 1). SH3 domains are interaction modules that typically bind to proline-rich motifs^60^. AlphaFold2 predictions of the three SH3 domains of CD2AP and CIN85, each in complex with an ALIX peptide containing R_745,_ highlight the importance of R_745_ which forms a key salt-bridge with a conserved glutamic acid within each SH3 domain, with additional acidic residues in the vicinity likely contributing favorably to the interaction (Fig. 4d, Supplementary Fig. 8 and 9). Similarly, R_757_ from an ALIX peptide establishes a salt-bridge with E_324_ in the unique SH3 domain of endophilin-A2, with E_320_ potentially enhancing the interaction (Fig. 4d, Supplementary Fig. 9).

Altogether, the novelty of our findings lies in demonstrating that the interactions between ALIX and CD2AP, CIN85 and endophilin-A2 are negatively regulated by arginine methylation of key residues within the proline-rich motif of ALIX, which are essential for SH3-mediated interactions with these proteins.

### The ability of ALIX to interact with CD2AP, CIN85 and endophilin-A2 is required for cytokinesis

To investigate the functional interaction between ALIX and these arginine methyl-sensitive partners, we generated a triple mutant of ALIX, ALIX-3K (R_745_K, R_757_K, R_767_K). Of note, we noticed that CD2AP, and, to a lesser extent endophilin-A2, were less expressed in cells transfected with GFP-ALIX compared to cells transfected with GFP (Fig. 5a and b) or GFP-ALIX-3K (Fig. 5a), suggesting that wild-type ALIX may promote their degradation. Co-immunoprecipitation experiments showed that ALIX-3K, in contrast to wild-type ALIX, was unable to associate with CD2AP and endophilin-A2, similar to when the arginine residues were methylated (see above Fig. 4), but retained the ability to interact with CEP55 and CARM1 (Fig. 5a). Co-expressing ALIX with CARM1 to increase ALIX methylation levels reduces or abolishes the binding of ALIX to CD2AP and endophilin-A2 (Fig. 5b). Conversely, an enzymatically inactive CARM1 mutant preserved the interaction (Fig. 5b). Together, these results indicate that either methylation of these arginine residues or substituting them with lysine leads to similar cellular consequences, specifically, the loss of SH3-mediated association of CD2AP and endophilin-A2 with ALIX.

**Figure 5.**
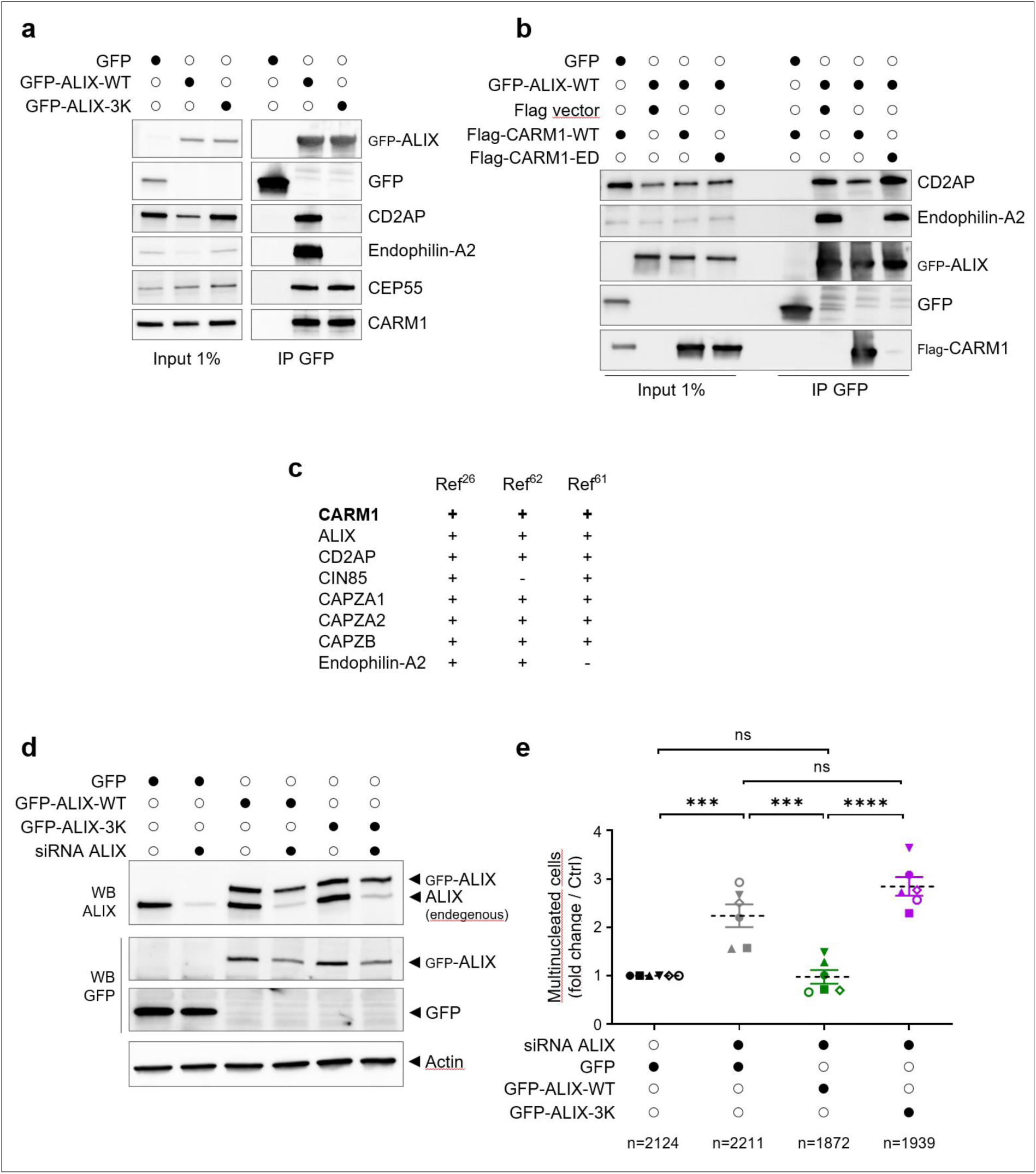
The ability of ALIX to interact with CD2AP, CIN85 and endophilin-A2 is required for cytokinesis. **(a)** HEK-293T ALIX knockout cells were transfected with the constructs as indicated. Protein extracts were immunoprecipitated (IP) using GFP-trap and the interactions with GFP-ALIX were assessed by western blot (*N* = 3). **(b)** HEK-293T ALIX knockout cells were co-transfected with GFP-ALIX-WT and with Flag-CARM1-WT or Flag-CARM1-ED (enzyme-dead mutant, CARM1-E_267_Q) or with control plasmids as indicated. IP using a GFP-trap was performed to assess the interactions between GFP-ALIX and CD2AP or endophilin-A2 by western blot (*N* = 2). **(c)** Table presenting the detection of various proteins of interest in purified midbodies. Data were retrieved from 3 studies that deciphered the proteome of midbodies^26,61,62^. **(d,e)** HeLa cells stably expressing GFP, GFP-ALIX-WT or GFP-ALIX-3K (ALIX siRNA-resistant) were transfected with ALIX siRNA to deplete endogenous ALIX. Western blots confirming the depletion of endogenous ALIX upon ALIX siRNA treatment. Actin was used as a loading control (d). Quantification of multinucleated cells in the four conditions (e). Error bars are mean ± SD. Six independent experiments were performed (mentioned by different symbols). The number of scored cells is > 250 per condition and per experiment (the total number of scored cells are indicated for each condition). ***p<0.001; ****p<0.0001; ns, not significant (Student’s t-test). Images shown are representative of two (b) three (a) or six (e) independent experiments.

ALIX is involved in various cellular processes requiring membrane remodeling, such as endocytosis, multivesicular body formation, virus budding, and cytokinesis. In this study, we focused on cytokinesis based on the analysis of midbody remnant proteomes published by us^26^ and others^61,62^, which showed the presence of not only ALIX, CD2AP, CIN85, endophilin-A2 and the capping proteins, but also CARM1 within this structure (Fig. 5c). Furthermore, the depletion of ALIX, CD2AP, CIN85, endophilin-A2 and the capping proteins has been reported to induce cytokinetic defects such as abscission delay and abscission failure leading to binucleation^36,37,63^. Therefore, we investigated whether CARM1-driven methylation of ALIX, which impairs association with CD2AP, CIN85, capping proteins, and endophilin-A2, affects the function of ALIX during the final step of cell division. We generated HeLa cell lines stably expressing GFP, siRNA-resistant GFP-ALIX-WT or GFP-ALIX-3K. GFP-ALIX-WT and GFP-ALIX-3K were expressed at levels comparable to those of endogenous ALIX (Fig. 5d). Depleting endogenous ALIX in GFP cells resulted in a 2.2-fold increase of multinucleated cells (Fig. 5d and e), in agreement with others^24,25^. These cytokinetic defects observed upon depletion of endogenous ALIX were fully rescued by expressing GFP-ALIX-WT (Fig. 5d and e). In contrast, no rescue was observed upon expressing GFP-ALIX-3K (Fig. 5d and e). These results indicate that the interaction mediated by R_745_, R_757_, R_767_ of ALIX with CD2AP, CIN85, capping proteins and endophilin-A2, without excluding the involvement of other partners, is required for ALIX’s function during cytokinesis.

## Discussion

The role of the arginine methyltransferase CARM1 as an enzyme involved in regulating gene expression is well established^13^. Here, we revealed a cytoplasmic function of CARM1 in regulating cell division through ALIX methylation. Our analysis of CARM1 interactomes across various cell lines identified the cytoplasmic protein ALIX as a main binding partner of CARM1, consistently ranking among the top five proteins in all CARM1 immunoprecipitates. We show that ALIX interacts with the catalytic domain of CARM1. Despite the catalytic domain being conserved across PRMTs, we found that ALIX specifically binds to CARM1 and not to the other PRMTs examined. Furthermore, we demonstrated that CARM1 associates with the intrinsically disordered PRD of ALIX, and methylates at least two arginine residues (R_745_ and R_757_) in cells and three arginine residues (R_745_, R_757_ and R_767_) *in vitro*. This result is consistent with the fact that arginine methylation often occurs in intrinsically disordered regions^5,51^. Moreover, we found that CARM1 associates with ALIX mutated at these arginine residues (R_745_K, R_757_K and R_767_K), implying that these amino acids are dispensable for CARM1-ALIX binding.

Protein methylation is considered to propagate signaling pathways by creating docking sites for methyl readers. Consistently, others have identified methyl readers for CARM1 and other protein methyltransferases using a similar peptide pull-down approach^64,65^. Instead, we identified proteins specifically associating with unmethylated arginine residues of ALIX. While only few reports have shown that CARM1-mediated arginine methylation can impair protein-protein interactions^66,67^, our findings underscore the importance of this regulatory mechanism and identified proteins whose interaction with ALIX are impaired through arginine methylation at R_745_ and R_757_ residues. This highlights the need for caution when utilizing “non-methylatable mutants” (arginine often substituted with lysine or alanine). While these mutants cannot undergo methylation, they can lead to misleading interpretations. Indeed, we would have incorrectly concluded that arginine methylation at R_745_ and R_757_ is required for interaction with CD2AP, CIN85 and endophilin-A2, based solely on the mutants’ inability to bind. The accurate conclusion was obtained using synthetic unmethylated and dimethylated peptides.

Remarkably, our unbiased pull-down approach identified CD2AP/CIN85 and endophilin-A2, previously reported to interact through their SH3 domains, with P_740_TPAP<u>R_745_</u> and P_755_A<u>R_757_</u>PPPP_761_ of ALIX, respectively^23,25,29,42^ (Supplementary Fig. 1). Mutation of two adjacent residues, including the CARM1-targeted arginine residue (underlined) in each of these motifs, impairs association of ALIX with CD2AP/CIN85 and endophilins^23,24^. In addition, substitution of R_745_ in an ALIX peptide with any other amino acid impairs binding to the SH3 domains of CD2AP *in vitro*^42^. In cells, we found that mutating either R_745_ or R_757_ to lysine impairs the interaction of ALIX with CD2AP and endophilin-A2, respectively. Crystal structures of the SH3 domains of CD2AP in complex with the PxPxP<u>R</u> motif of RIN3 (corresponding to P_740_TPAP<u>R_745_</u> in ALIX) have been resolved, revealing that the critical arginine residue (underlined in the motif) interacts with an acidic patch in the SH3 domains of CD2AP^42,68^. The structural models generated by AlphaFold2 between ALIX fragments and the SH3 domains of CD2AP, CIN85 and endophilin-A2 converge to a similar configuration where the arginine residues in the poly-proline motif, R_745_ and R_757_ also engage with acidic residues to form stabilizing salt-bridges. Hence, the arginine residues methylated by CARM1, are key residues interacting with conserved acidic residues in SH3 domains, and the association is impaired when they are mutated or methylated (Fig. 6). The novelty of our study is the demonstration that the interactions between ALIX and CD2AP, CIN85 and endophilin-A2 are negatively regulated by arginine methylation, by impairing binding between proline-rich and SH3 domains. Thus far, only one case of arginine methylation in a proline rich motif of HIV-1 Nef has been reported to modulate its interaction with the SH3 domain of Fyn^67^. Inspection of the structural models and of the distribution of atomic contacts made by the proline residues surrounding R_757_, suggest that the motif for binding to endophilin-A2 could rather be Px<u>R</u>PxxP. Interestingly, dozens of additional proteins contain a PxPxP<u>R</u> motif, which represents the minimum sequence for binding to SH3 domains of CD2AP and CIN85^42,69^. Our work thus raises the possibility that arginine methylation represents a post-translational modification that may be widely used to modulate the interaction between proline-rich motifs and SH3 domain-containing proteins.

**Figure 6.**
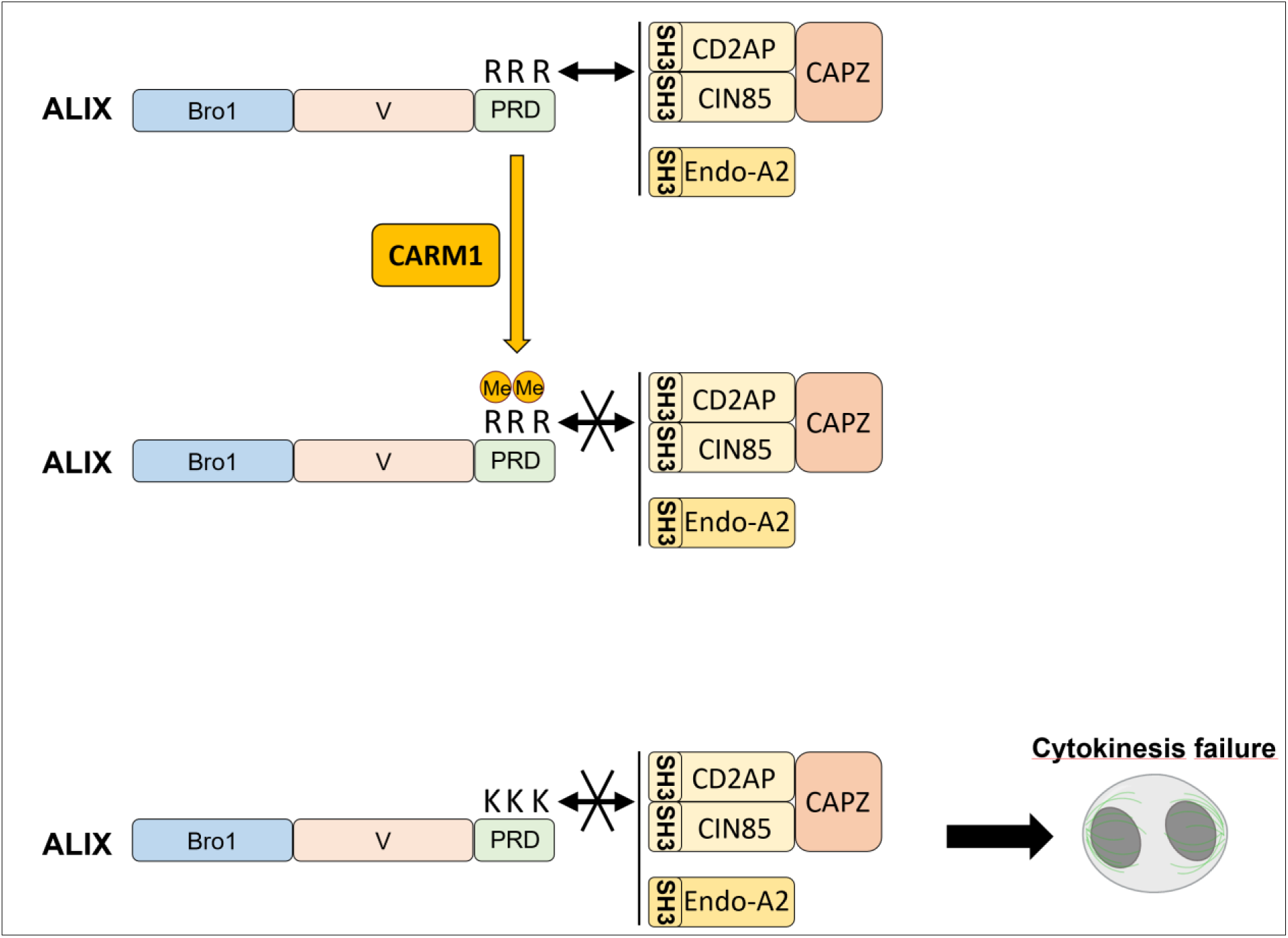
Working model: CARM1-dependent methylation of the proline-rich domain of ALIX impairs SH3-mediated interactions with CD2AP, CIN85 and endophilin-A2, whose association with ALIX are critical for successful cytokinesis. Double ended arrows represent direct interactions.

Our previous quantitative analysis, as well as publicly available proteomes of purified midbodies^26,61,62^, identified the presence of CARM1, in addition to proteins known to regulate cytokinesis, such as ALIX, CD2AP, CIN85, endophilin-A2 and capping proteins. These findings suggest that CARM1 could regulate cytokinesis through arginine methylation of target proteins at the midbody. ALIX is a master regulator of the completion of cytokinesis by recruiting the ESCRT-III machinery (via its Bro1 domain, Supplementary Fig. 1) to the midbody to initiate its polymerization. Furthermore, ALIX stabilizes the ESCRT-III filaments with the plasma membrane by interacting with syntenin/syndecan-4 (Supplementary Fig. 1), thus ensuring the stable association of ESCRT-III at the abscission site^16^. We show that the use of an ALIX mutant demonstrated the requirement of ALIX to interact with other partners (CD2AP, CIN85, capping proteins, endophilin-A2, without excluding additional partners) for successful cytokinesis. Of note, cells expressing ALIX mutants, which are incapable of binding to these proteins, have been reported to partially or fully rescue cytokinetic defects due to the loss of endogenous ALIX^23,25^. This apparent discrepancy between these results and ours could be explained by (i) the use of different HeLa cell lines displaying variable percentages of multinucleated cells upon ALIX depletion^23,25^, (ii) the use of different methods to assess cytokinetic defects (quantification of multinucleated cells in our study versus quantification of polyploid (8n) cells^25^, and/or more importantly, (iii) the use of an ALIX mutant defective for both CD2AP/CIN85 and endophilin-A2 binding sites in our study compared to the use of single-binding site mutants^23,25^.

The silencing of CD2AP, endophilin-A2 or capping proteins results in cytokinetic defects with uncharacterized molecular mechanisms^37,63^, except for the depletion of actin-capping proteins resulting in an accumulation of actin filaments (F-actin) in the intercellular bridge^36^. ALIX interacts with F-actin and related proteins^70-74^; however, whether ALIX controls F-actin cytoskeleton within the intercellular bridge is unknown. CD2AP and CIN85 also interact with actin-associated proteins, such as anillin, septins, and capping proteins, which are essential for regulating actin dynamics and contractility required for intercellular bridge maturation^38,63,75^. Capping proteins prevent both polymerization and depolymerization of F-actin^76^, a process inhibited when they interact with CD2AP^75,77^. Actin-capping proteins have recently been shown to bind to the Bro1 domain of ALIX^35^. In our study, we identified the PRD domain of ALIX as an additional binding site for these proteins. Given that capping proteins have been reported to interact with CD2AP/CIN85^56-59^, we hypothesize that CD2AP/CIN85 may serve as a bridge between ALIX to capping proteins, forming a complex that may regulate F-actin dynamics during cytokinesis. CARM1-mediated methylation of ALIX could therefore regulate actin remodeling in the intercellular bridge; however, this hypothesis warrants further investigation. In addition, our results suggest that ALIX could regulate cytokinesis through its association with endophilin-A2. Although the function of endophilin-A2 in this process is uncharacterized, it has been shown to control vesicular trafficking in other contexts than cytokinesis. Vesicular trafficking delivers key proteins such as ALIX, ESCRT-III, and enzymes catalyzing F-actin-disassembly^21,78-80^ to the intercellular bridge and to the midbody, facilitating the final steps of cytokinesis. As ALIX cooperates with endophilin-A2 during endocytosis, they could also control vesicular trafficking required for the final steps of cytokinesis, and this process could be regulated by CARM1-mediated methylation of ALIX.

In summary, our study demonstrates that the methylation of ALIX by CARM1 impairs SH3-dependent interactions with key cytokinetic proteins, CD2AP, CIN85, and endophilin-A2, and that these interactions are crucial for ALIX function in cytokinesis (Fig. 6). This supports the notion that protein methylation is a major PTM governing the last step of cell division. Indeed, methylation of the ESCRT-III subunit CHMP2B on lysine residues by the lysine methyltransferase SMYD2 regulates cytokinesis possibly by modifying CHMP2B interactions with membranes, as recently reported^81^. It is remarkable that several ESCRT or ESCRT-associated components are regulated by different methylation modifications (lysine methylation for CHMP2B and arginine methylation for ALIX) during cytokinesis. CD2AP, CIN85, and endophilin-A2 have well-established functions outside cell division. Therefore, our study raises the possibility that CARM1-dependent methylation of ALIX regulates key cellular functions beyond cell division, such as multivesicular body formation, extracellular vesicle biogenesis, virus budding, membrane repair or endocytosis.

## Methods

### Cell lines and cell culture

BT-549, HEK-293T and HeLa CCL-2 cells were purchased from the American Type Culture Collection (ATCC). MDA-MB-231 cells were a kind gift from Mina Bissell (Berkeley Lab, University California). HEK-293T ALIX knockout (KO) were obtained from Rémy Sadoul. BT-549 cells were maintained in RPMI-1640 glutamax supplemented with 10% fetal bovine serum (FBS), 1% penicillin-streptomycin (PS), 1.5 g/L Sodium bicarbonate, 10 mM Hepes, 1 mM Sodium pyruvate. HEK-293T, HeLa, HeLa GFP, HeLa GFP-ALIX-WT and HeLa GFP-ALIX-3K cells were maintained in DMEM glutamax (Gibco, Lifetechnologies) supplemented with 10% FBS and 1% PS. Cell lines were authenticated in 2021^82^ by short tandem repeats and tested for mycoplasma using MycoAlert Mycoplasma Detection Kit (Lonza Biosciences). HeLa cell lines stably expressing GFP, GFP-ALIX and GFP-ALIX-3K were generated using transposition and the iON strategy^83^. HeLa cells were seeded in 6-well plates and transfected the following day using Xtremegene-9 DNA reagent (Roche) for 48 h with a mixture of 0.2 µg iON-HypBASE-transposase plasmid + 1 µg of either iON-CAG-GFP, iON-CAG-GFP-ALIX-siRNA-resistant or iON-CAG-GFP-ALIX-3K-siRNA-resistant plasmids. Cells were then selected by FACS to obtain a bulk population of GFP-positive cells expressing the fusion proteins at levels comparable to those of endogenous ALIX. Plasmids and siRNAs are reported in Supplementary material Tables 1 and 2.

### Inhibitors

EZM2302 (Probechem Biochemicals, PC-61030), TP064N (Structural Genomics Consortium) and TP064 (Tocris, 6008) were resuspended in DMSO. HeLa ATCC and BT-549 cells were treated with the indicated concentrations of CARM1 inhibitors (EZM2302 and TP064) or negative control (TP064N or 0.01% DMSO) for 48 h.

### Transfections with siRNAs

For silencing experiments, HeLa cells were transfected at the time of seeding and again 24 h later with 20 nM siRNAs using Lipofectamine RNAimax reagent (Invitrogen) for a total transfection duration of 48 h. Stable HeLa cell lines and MDA-MB-231 cells were transfected with 20 nM siRNAs for 48 h using Interferin (PolyPlus) or Lipofectamine RNAimax reagent (Invitrogen), following the manufacturer’s instructions. For plasmid transfection, HEK-293T and HEK-293T ALIX-KO were transfected using Xtremegene-HP DNA reagent (Roche) for 48 h, following the manufacturer’s instructions.

### Plasmids

pcDNA3.1(+) plasmid was from Lifetechnologies (#V79020). Flag-CARM1-FL, Flag-CARM1-FL-R_168_A, Flag-CARM1-FL-E_267_Q and Flag-PRMT5 were synthesized and cloned into pcDNA3.1(+) by Genewiz. Flag-CARM1 del15 was generated from Flag-CARM1-FL using site-directed mutagenesis (Quickchange II XL kit, Agilent). Flag-PRMT1 was generated by replacing PRMT5 by PRMT1 in the Flag-PRMT5 vector using NEBuilder HiFi DNA Assembly Cloning Kit (New England BioLabs, E5520). Flag-PRMT3 was from Addgene (#164695). Flag-PRMT2 was generated from pGex-6P-1-GST-PRMT2 (Addgene, #34650). pEGFP-C1 was a kind gift from Ludger Johannes (Institut Curie). GFP-ALIX-FL, mcherry-ALIX-FL, mcherry-ALIX-Bro1, mcherry-ALIX-ΔBro1, iON-CAG-MCS and iON-HypBASE-transposase were from Arnaud Echard’s laboratory. The truncated forms of Flag-CARM1, mcherry-ALIX and GFP-ALIX were obtained by standard cloning or mutagenesis (Quickchange II XL kit, Agilent) strategies using the primers listed in Supplementary material Table 1. GFP-ALIX-3K (R_745_K, R_757_K, R_767_K) was generated by site-directed mutagenesis (Quickchange II XL kit, Agilent). siRNA-resistant versions of GFP-ALIX and GFP-ALIX-3K were obtained by mutating six nucleotides in the ALIX siRNA-targeting sequence (Genewiz). To generate iON-CAG-GFP, iON-CAG-GFP-ALIX-WT and iON-CAG-GFP-ALIX-3K, the GFP, GFP-ALIX and GFP-ALIX-3K coding sequences were amplified by PCR as described^83^ from peGFP-C1, GFP-ALIX-siRNA-resistant and GFP-ALIX-3K-siRNA-resistant plasmids, respectively, and inserted into iON-CAG-MCS vector using NEBuilder HiFi DNA Assembly Cloning Kit (New England BioLabs, E5520). pET-11-GB1-His6-TEV-ALIX-PRD_703-800_-Strep was obtained from Addgene (#141344)^84^ and was used to generate pET-11-GB1-His6-TEV-ALIX-PRD_703-800_-Strep-R_745_K-R_757_K (ALIX-PRD_703-800_-2K) and pET-11-GB1-His6-TEV-ALIX-PRD_703-800_-Strep-R_745_K-R_757_K-R_767_K (ALIX-PRD_703-800_-3K) by site-directed mutagenesis (Quickchange II XL kit, Agilent). GST from pGEX-4T-1 was cloned into the pFastBac vector. CARM1-FL sequence from Flag-CARM1-FL was cloned into the pFastBac-GST vector. All constructs were verified by DNA sequencing (Eurofins genomics, France) and encode for human proteins. Cloning and mutagenesis primers are listed in Supplementary material Table 1.

### Western blot

Total proteins were extracted in Laemmli buffer (50 mM Tris pH 6.8, 2% SDS, 5% glycerol, 2 mM dithiothreitol (DTT), 2.5 mM ethylenediaminetetraacetic acid (EDTA), 2.5 mM ethylene glycol tetraacetic acid (EGTA), 4 mM sodium orthovanadate, 20 mM sodium fluoride, phosphatase and protease inhibitors) and heated for 10 min at 100°C. Protein concentrations were determined using the Pierce BCA assay kit (ThermoFisher) and Bromophenol blue (1%) was added to protein lysates. Proteins (20 µg) were loaded on Mini-PROTEAN TGX (4-15%) precast gels (Bio-Rad) for migration, and transferred onto nitrocellulose membranes (Trans-Blot Turbo Mini 0.2 μm, Bio-Rad). Protein loading and transfer were assessed by stain-free imaging technology (Bio-Rad Laboratories). The membranes were saturated with 5% bovine serum albumin (BSA) (Sigma Aldrich) in Tris-Buffered Saline (Interchim) containing 0.1% Tween-20 (TBST) (Amresco) for 1 h and incubated with primary antibodies overnight at 4°C. After three washes in TBST, the membranes were hybridized with the secondary antibody coupled to peroxidase for 1 h at room temperature (RT). The antibodies (listed in Supplementary material Table 3) were diluted in 5% BSA + TBST. The membranes were washed three times with TBST and immune complexes were revealed by enhanced chemiluminescence (SuperSignal West Pico PLUS Chemiluminescent Substrate, ThermoFisher) and imaged using a ChemiDoc™ XRS+ System (Bio-Rad Laboratories).

### Immunoprecipitation

Cells were washed with ice-cold PBS, lysed by adding cold complete lysis buffer (50 mM Tris pH 7.4, 100 mM NaCl, 1 mM EDTA, 1 mM EGTA, 0.1 to 1% Nonidet P-40 (NP-40), 1 mM DTT, 10% glycerol, phosphatase and protease inhibitors), detachedusing a cell scraper and collected into microcentrifuge tubes, and left under rotation (40 rpm) at 4°C for 30 min before centrifugation (4°C, 15 min, 13,200 rpm). Protein concentration was determined using the Pierce BCA assay kit (ThermoFisher). Protein lysates were diluted at 1 mg/mL in a final volume of 1 mL and incubated overnight at 4°C with 2 µg of either the antibody of interest or an isotype control antibody (e.g., rabbit IgG or mouse IgG) under rotation at 40 rpm. Pierce Protein G agarose beads (Life technologies) were washed three times in lysis buffer, and 20 μL of beads were added to each protein lysate, followed by incubation for 1 h at 4°C. The samples were centrifuged for 1 min at 13,200 rpm at 4°C and washed three times in 1 mL lysis buffer. The immunoprecipitates were resuspended in 20 µL Laemmli buffer, heated for 10 min at 100°C and analyzed by western blot. Flag- and GFP-tagged proteins were precipitated using ChromoTek (Flag or GFP)-Trap™ Agarose (ProteinTech). The beads were washed three times with lysis buffer, and 20 μL of the washed beads were added to 0.5 mg of protein lysate (0.5 mg/mL) and incubated for 2 hours at 4°C under rotation (40 rpm). Following incubation, the beads were centrifuged and washed as described above. The antibodies used are listed in Supplementary material Table 3.

### Proteomics Sample Preparation for MS analysis

To identify CARM1, Flag-CARM1-FL/Δ15 and ALIX interactomes (hereafter referred to as “CARM1 and ALIX interactomes”) and ALIX post-translational modifications hereafter referred to as “ALIX PTM”), proteins were immunoprecipitated as described above. After three washes in lysis buffer, beads were resuspended in 100 μL of 25 mM NH4HCO3 or in 100 μL of 25 mM NH4HCO3 plus 10 mM CaCl_2_ for CARM1 and ALIX interactomes and ALIX PTM, respectively. On-bead digestion was done by adding 0.2 μg of Tryspsin/LysC (Promega) to CARM1 and ALIX interactome beads and by adding 0.4 μg of LysArginase (Sigma EMS0008) to ALIX PTM beads for 1 h at 37 °C. The resulting peptide mixtures were then loaded onto homemade C18 StageTips packed with AttractSPE™ Disks Bio C18 (Affinisep™ SPE-Disks-Bio-C18-100.47.20) for desalting. Peptides were eluted using 40/60 CH3CN/H2O + 0.1% formic acid, vacuum concentrated to dryness and reconstituted in injection buffer in 0.3% Trifluoroacetic acid (TFA) before Liquid chromatography-tandem mass spectrometry (LC-MS/MS) analysis. CARM1 and ALIX interactomes were analyzed by data dependent acquisition (DDA) and ALIX PTM also in parallel reaction monitoring (PRM) mode.

### LC-MS/MS analysis

Online chromatography was performed with an RSLCnano system (Ultimate 3000, Thermo Scientific) coupled to a Orbitrap Exploris 480 (Thermo Scientific). Peptides were trapped on a C18 column (75 μm inner diameter × 2 cm; nanoViper Acclaim PepMapTM 100, Thermo Scientific) with buffer A (2/98 CH_3_CN/H_2_O in 0.1% formic acid) at a flow rate of 3.0 µL/min over 4 min. Separation was performed on a 50 cm x 75 μm C18 column (nanoViper Acclaim PepMapTM RSLC, 2 μm, 100 Å, Thermo Scientific) regulated to a temperature of 40°C for CARM1 and ALIX interactomes and 50°C for ALIX PTM, respectively, and with a linear gradient of 3% to 29% buffer B (100% CH_3_CN in 0.1% formic acid) for CARM1 and ALIX interactomes and 2% to 30% buffer B for ALIX PTM, at a flow rate of 300 nL/min over 91 min. DDA was performed in the ultrahigh-field Orbitrap mass analyzer in ranges m/z 375–1500 with a resolution of 120 000 at m/z 200. In PRM mode, for ALIX PTM analysis, MS2 scan parameters were set to select the m/z ratio of ALIX peptides from the inclusion list (Table A) generated from the peptides obtained from previous DDA. For the CARM1 and ALIX interactomes, background signal was accounted for by subtracting the number of peptides detected in the control IP condition from those detected in the CARM1 or ALIX IP conditions. Proteins were then ranked from highest to lowest based on the resulting peptide counts.

**Table A:**
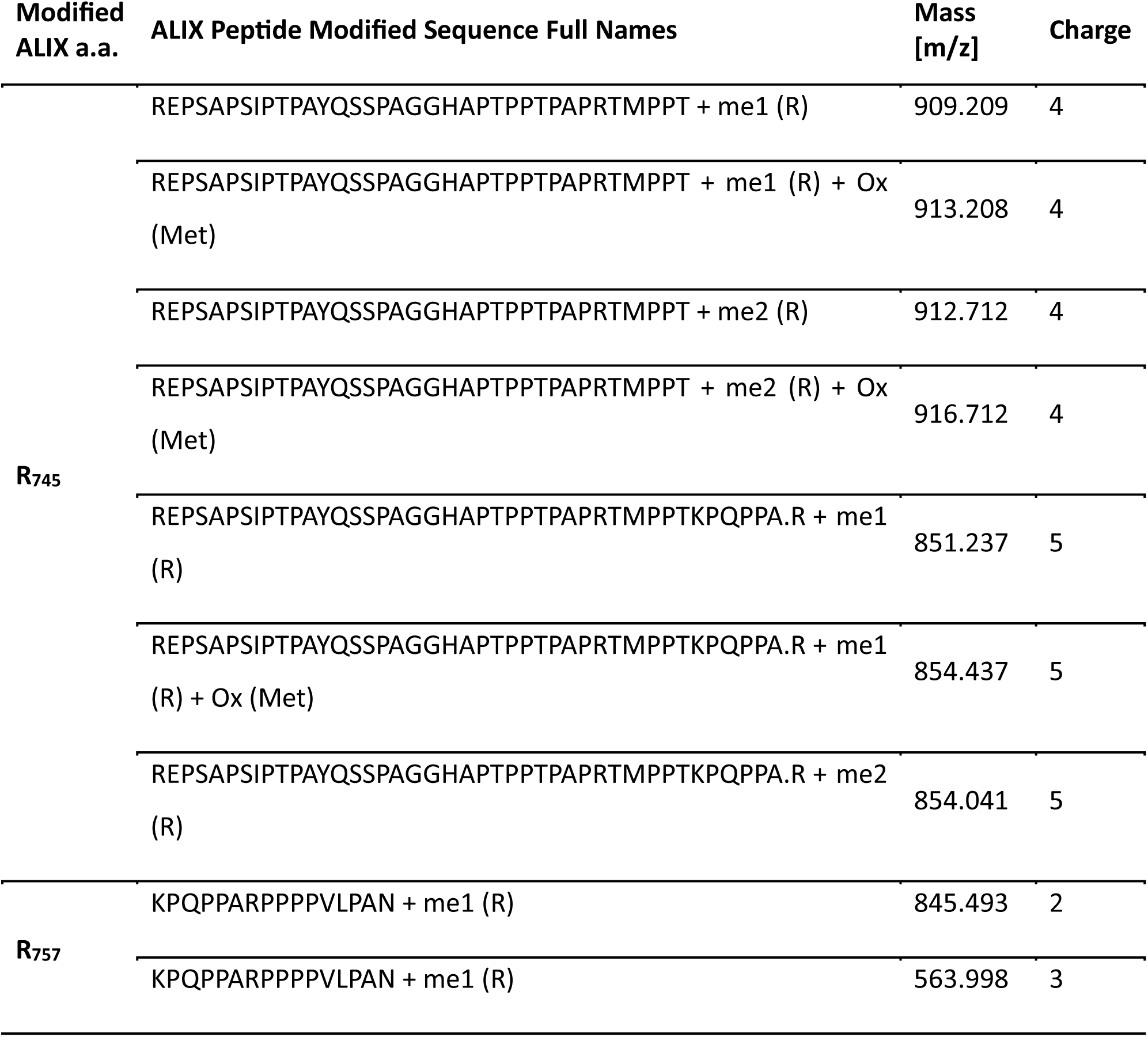
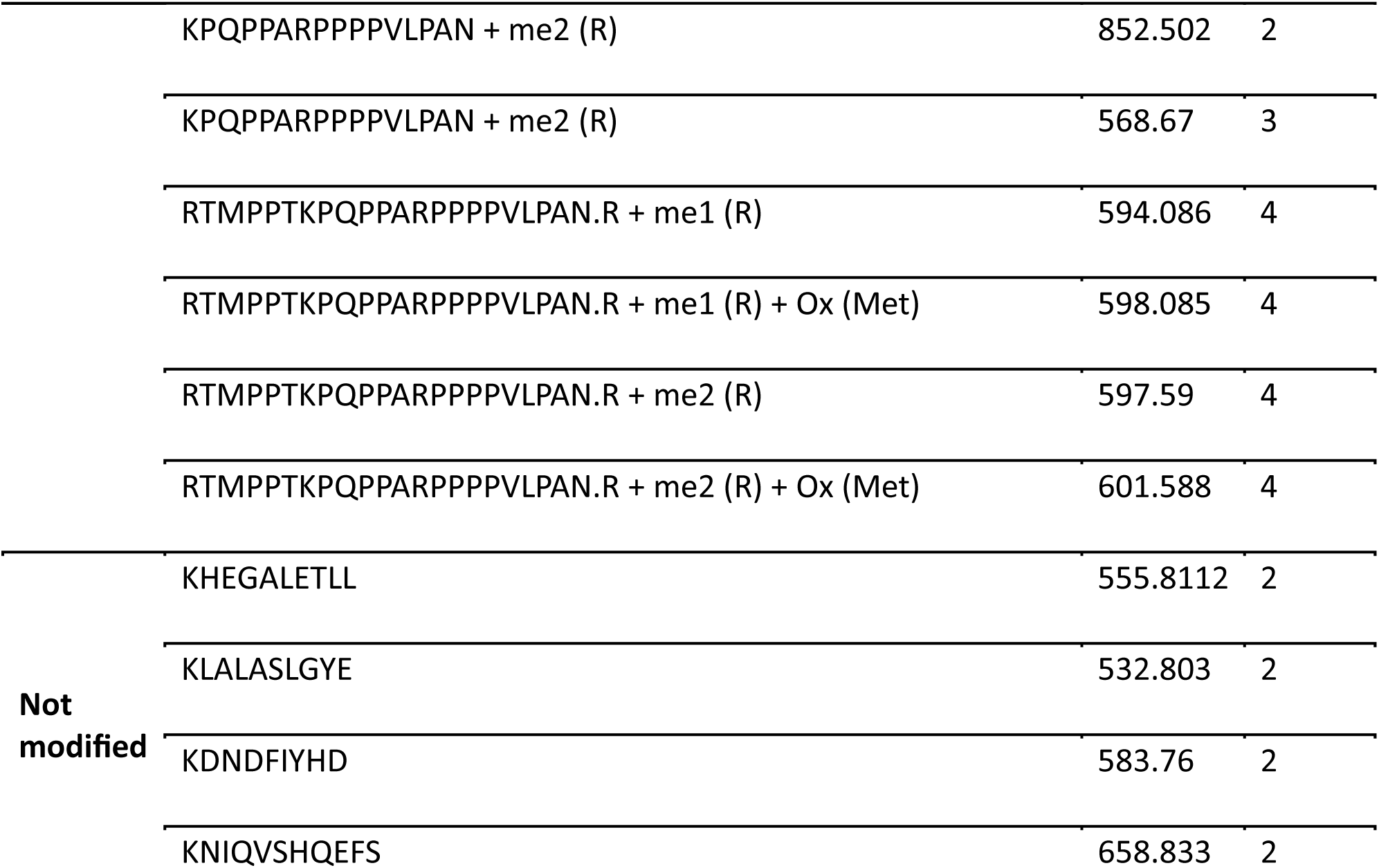
Inclusion list generated from the ALIX peptides obtained from previous DDA used to set the M2 scan parameters and select the m/z ratio of ALIX peptides for in vivo ALIX PTM analysis in PRM mode.

### Data processing of MS files

For the protein identification of CARM1 and ALIX interactomes, the data were searched against the Homo sapiens (UP000005640) UniProt database using Sequest HT through Proteome Discoverer (version 2.4). Enzyme specificity was set to Trypsin (N-terminal cleavage of lysine and arginine) and a maximum of two-missed cleavage sites were allowed. Oxidized methionine, Met-loss, Met-loss-Acetyl and N-terminal acetylation were set as variable modifications and carbamidomethylation of cysteins was set as fixed modification. Maximum allowed mass deviation was set to 10 ppm for monoisotopic precursor ions and 0.02 Da for MS/MS peaks. The resulting files were further processed using myProMS v3.10^85^ (https://github.com/bioinfo-pf-curie/myproms). FDR calculation used Percolator^86^ and was set to 1% at the peptide level for the whole study.

For ALIX PTM quantification, raw files were processed using Skyline (version 22.2.0.527) MacCoss Lab Software (Seattle, WA, https://skyline.ms/project/home/software/Skyline/begin.view) to generate the extracted-ion chromatograms and peak integration. Since we only have one protein, mean-scaled normalization was applied on non-modified peptides, considering the experimental repeats. The coefficients of normalization were then applied to the modifications. To evaluate the statistical significance of the change in protein abundance, a linear model (adjusted on peptides, biological replicates and experimental repeats) was performed, and a T-test was applied on the fold change estimate. The p-values were then corrected for multiple testing using the Benjamini-Hochberg procedure.

### Quantification of multinucleated cells

After siRNA transfection, cells seeded on glass coverslips were fixed with 4% paraformaldehyde (PFA) at RT for 15 min. Cells were washed twice in PBS and permeabilized with 0.1% Triton-X-100 in PBS for 10 min at RT. Nuclei were stained with 0.1 μg/mL Dapi. Final washes were performed using PBS + 1% BSA and PBS before mounting the coverslips with Mowiol 4-88 (Sigma-Aldrich) solution. Images were acquired using an Upright Epifluorescence widefield with Apotome microscope (Zeiss) at 40X (numerical aperture = 1.3) or 63X (numerical aperture = 1.4) and a CoolSnap HQ2 camera.

### Recombinant protein production

GST-CARM1 recombinant protein was produced using the modified Bac-to-Bac Baculovirus expression system (Invitrogen, 10359-016). Briefly, the GST-CARM1-FL was cloned in pFastbac/EGFP vector and transformed into *Escherichia coli* (*E. coli*) DH10Bac-LL to generate a recombinant bacmid-LL. Sf9 insect cells were grown in Insect-XPRESS (Lonza 2-730Q) at 28°C under rotation (120 rpm) and transfected with a mixture of the bacmid DNA and PEI-MAX 40 (polyscience 24765 (100)) transfectant. The medium containing the freshly produced baculoviruses was collected 5 days post-transfection, controlled by the fluorescence of EGFP and used to infect a large culture of Sf9 cells for viral amplification. To produce the recombinant GST-CARM1, Sf9 cells were infected with the recombinant baculovirus at a multiplicity of infection (MOI) of 10. After 72 h, the cells were harvested and incubated for 1 h at 4°C in buffer C (50 mM Tris pH 8, 300 mM NaCl, 5 mM β-mercaptoethanol, 10% glycerol) supplemented with 0.5% Triton X-100 and a complete protease inhibitor cocktail (Roche). The clarified cell lysate was centrifuged at 22,000 rpm for 1 h to remove debris. The supernatant was then loaded onto a 1 mL Glutathione Sepharose beads column (Cytiva) using an Akta-pure system, which had been pre-equilibrated with buffer C at 4°C. Recombinant proteins were eluted from the column with buffer C supplemented with 50 mM reduced glutathione. The eluted fractions containing the GST-CARM1 protein were concentrated and subjected to size exclusion chromatography on a Superdex 200 10/300 column, also equilibrated with buffer C. The fractions containing the recombinant protein were pooled, and the final protein concentration was determined by measuring the absorbance at 280 nm.

Purified recombinant protein of human Flag-ALIX produced in FreeStyle HEK-293T cells was provided by Christine Chatellard and Rémy Sadoul. Recombinant human histone H3 (H3.1) was from New England Biolabs.

GB1-His6-ALIX-PRD_703-800_-Strep (ALIX-PRD_703-800_-WT), GB1-His6-ALIX-PRD_703-800_-R_745_K-R_757_K-Strep (ALIX-PRD_703-800_-2K) and GB1-His6-ALIX-PRD_703-800_-R_745_K-R_757_K-R_767_K-Strep (ALIX-PRD_703-800_-3K) recombinant proteins were produced in bacteria. The pET-11-GB1-His6-TEV-ALIX-PRD_703-800_-Strep, GB1-His-ALIX-PRD_703-800_-Strep-2K and GB1-His-ALIX-PRD_703-800_-Strep-3K vectors were transformed into BL21(DE3) *E. coli*. Protein production was induced by adding 0.1 mM Isopropyl β-d-1-thiogalactopyranoside (IPTG) to the bacterial suspension at an optical density between 0.6 and 0.8. Cells were cultured for 24 h at 16°C under rotation and lysed with 50 mM Tris pH=8, 1% Triton X-100, 500 mM NaCl, 50 mM EDTA, 5% glycerol, 1 µg/mL leupeptin, 1 µg/mL aprotinine, 1 mM PMSF, protease inhibitors (Roche), 10 mg/mL lysozyme, 3 units of benzonase, and 2 mM MgCl_2_. The recombinant proteins were purified using a Strep-Tactin® Sepharose® resin (IBA-Lifesciences, 2-1201-002) according to the manufacturer’s protocol.

### *In vitro* methylation assay

The *in vitro* methyltransferase reaction was carried out for 1.5 h at 30°C in methylation buffer (50 mM Tris pH=7.2, 50 mM NaCl, 1 mM DTT) containing 1 µg of recombinant proteins (GST-CARM1, Flag-ALIX, ALIX-PRD_703-800_-WT, ALIX-PRD_703-800_-2K or ALIX-PRD_703-800_-3K) and 0.5 μCi of ^3^H-S-adenosyl-L-methionine (^3^H-SAM, PerkinElmer) in a final volume of 20 µL. The reaction was stopped by the addition of 4X Laemmli buffer and heating the samples at 100°C for 10 min prior to loading on a Mini-PROTEAN TGX (4-15%) precast gels (Bio-Rad). Proteins were then transferred onto a PVDF membrane using a Trans-Blot SD Semi-Dry Transfer Cell (Bio-Rad) and stained with Coomassie blue. The membrane was placed in contact with a Phospho-screen BAS-IP TR 2025 E Tritium (Dutscher) for 48 h at RT and radioactive signal was visualised using a Typhoon™ laser-scanner (Cytiva).

### Peptide pull down assays and SILAC-based quantitative proteomic analyses

ALIX peptides (Supplementary material Table 4) were synthesized by CASLO ApS (Denmark). For each proteomic SILAC peptide pull-down, 10 µL of slurry Streptavidine T1 Dynabeads™ MyOne™ (Invitrogen) were saturated with 50 µg of specific biotinylated peptides in lysis/washing buffer (50 mM Tris pH 8, 150 mM NaCl, 1 % NP40, complete protease inhibitors (Roche)) for 2 h at 4°C under rotation. Next, beads were washed in the lysis/washing buffer and incubated for 4 h at 4°C under rotation with 1 mg of MDA-MB-231 total extract prepared from cells cultivated in either normal isotope amino acids culture condition (‘Light”, L-Arginine HCl unlabeled, L-Lysine HCl unlabeled. Silantes) or using modified isotope amino acids culture condition (‘Heavy’, L-Arginine HCl U-^13^C, U-^15^N, L-Lysine HCl U-^13^C, U-^15^N. Silantes). A two-way experiment was performed, the ‘forward’ condition combining ALIX-me0 peptide with light extract and ALIX-me2a peptide with heavy extract, the ‘reverse’ condition being a swap of the peptides and extracts. Beads of each pair of peptide pulldown were then pooled together, and extracts were resuspended in Laemmli buffer. Eluted proteins were stacked on top of a 4%– 12% NuPAGE gel (Invitrogen). After staining with R-250 Coomassie Blue (Biorad), proteins were digested in-gel using modified trypsin (sequencing grade, Promega) as described in https://www.ncbi.nlm.nih.gov/pmc/articles/PMC10830559/ - CR58^87^. The resulting peptides were analyzed by online nano liquid chromatography coupled to MS/MS (UHPLC Vanquish Neo coupled to Orbitrap Ascend Tribrid, Thermo Fisher Scientific) using a 60 min gradient. For this purpose, the peptides were sampled on a pre-column (300 μm × 5 mm PepMap C18, Thermo Scientific) and separated in a 75 μm × 250 mm C18 column (Aurora Generation 3, 1.7 μm, IonOpticks). The MS and MS/MS data were acquired by Xcalibur (Thermo Fisher Scientific). Peptides and proteins were identified and quantified using MaxQuant^88^ (2.4.2.0) using the Uniprot database (Homo sapiens taxonomy, 20240404 download) and the frequently observed contaminant database embedded in MaxQuant. Trypsin was chosen as the enzyme and 2 missed cleavages were allowed. Peptide modifications allowed during the search were: carbamidomethylation (C, fixed), acetyl (Protein N-ter, variable) and oxidation (M, variable). The minimum peptide length and the minimum number of razor peptides were respectively set to six amino acids and one peptide. Maximum false discovery rates — calculated by employing a reverse database strategy — were set to 0.01 at peptide-spectrum-match and protein levels. Quantification of SILAC ratios was performed using default settings. Proteins identified as outliers in both experiments are assigned as significant interactors.

### Peptide pull-down assays for western blot

Streptavidin Sepharose beads (20 µL) (GE Healthcare) were saturated with 50 µg of ALIX biotinylated peptides in a reaction buffer (50 mM Tris pH=7.4, 100 mM NaCl, 1 mM EDTA, 1 mM EGTA, 0.1% NP40, 1 mM DTT, 10% glycerol) for 2 h at 4°C under rotation (40 rpm). Then, the beads were washed three times with 1 mL of the same buffer. The beads were incubated overnight at 4°C under rotation (40 rpm) with 0.5 mg of total protein lysates from MDA-MB-231 or HeLa ATCC cells (0.5 mg/mL), which had been lysed in lysis buffer (reaction buffer supplemented with phosphatase and protease inhibitors). Beads were washed three times in 1 mL of reaction buffer. After removal of the final wash, the bound proteins were incubated at 100°C for 10 min in 25 µL of Laemmli buffer and analysed by western blot.

### Structural modeling

The structural modeling of protein complexes was performed using AlphaFold2^89^. The concatenated multiple sequence alignments of delimitated partners were generated using MMSeqs2^90^ and submitted as inputs to the ColabFold v1.5.2^91^ implementation of AlphaFold2 with the Multimer v2.3 model parameters^92^ with 3 recycles and 5 generated models. Four scores were provided by AlphaFold2 to rate the quality of the models, the pLDDT, the pTMscore, the ipTMscore and the model confidence score (weighted combination of pTM and ipTM scores with a 20:80 ratio). The scores obtained for all the generated models are reported Supplementary Figure 9.

### Data availability

The MS-based proteomics data have been deposited to the ProteomeXchange Consortium via the PRIDE^93^ partner repository and will be available once the manuscript will be published.

## Acknowledgements

This work was supported by the Institut Curie and by the European Union, EVCA Twining Project (Horizon GA n° 101079264) to TD and Institut Pasteur, CNRS, FRM (Recherche soutenue par la FRM EQU202103012627) and the Agence Nationale pour la Recherche (ANR 20 CE011 0014 SepScort) to AE. SS was funded by the European Union’s Horizon 2020 Research and Innovation Program (Marie Skłodowska-Curie grant agreement No 666003). SR was supported by the “Institut Curie EuReCa PhD Program”, co-funded by the European Union’s Horizon 2020 research and innovation program (Marie Skłodowska-Curie Actions-grant agreement No 847718), and by the Fondation ARC pour la recherche sur le cancer. RDa was financed by the French Embassy of Lebanon and the Lebanese University (Safar Volet 1). RDi was supported by Fondation ARC pour la recherche sur le cancer (ARCPOST-DOC2023080006931). JT was supported by the Pasteur-Paris University (PPU) international PhD program. We gratefully acknowledge Chloé Guedj, Mathieu Cortes and the Cell and Tissue Imaging Platform (PICT-IBiSA) at Institut Curie, member of the French National Research Infrastructure France-BioImaging (ANR-10-INBS-04). YC acknowledges the support by Agence Nationale de la Recherche under projects ProFI (Proteomics French Infrastructure, ANR-10-INBS-08) and GRAL, a program from the Chemistry Biology Health (CBH) Graduate School of University Grenoble Alpes (ANR-17-EURE-0003). We are grateful to the Institut Curie’s Cytometry Platform for its technical help and scientific discussion. We thank Mark Bedford (MD Anderson Cancer Center), Ludger Johannes (Institut Curie) and Mina Bissell (Lawrence Berkeley National Laboratory) for providing reagents. We thank Patrick Poullet (bioinformatics platform, Institut Curie U900) for the continuous development of myProMS and Michael Richard (bioinformatics platform, Institut Curie U900) for the statistical analyses of mass spectrometry data generated at Institut Curie. We thank Kardelen Gökçen for performing experiments not included in this manuscript, and Michel Wassef (Institut Curie) for advice. We thank Sergio Roman-Roman (Institut Curie), Olivier Destaing (University Grenoble Alpes), Guillaume Montagnac (Gustave Roussy Institute), Jocelyn Côté (University of Ottawa), Wei Xu (University of Wisconsin-Madison) and all members of the groups of Thierry Dubois, Arnaud Echard, Philippe Chavrier (Institut Curie), Jonathan Weitzman and Souhila Medjkane (Paris Cité University) for helpful discussions. The BIOI2 platform resources at the I2BC is acknowledged for the implementation of the AlphaFold2 service.

## Author contributions

SH and SS performed most of the experiments and their analysis, with support from TD. SS characterized the CARM1 and ALIX interactomes, while SR characterized the endogenous CARM1 isoform interactomes. SH, SS, and CV validated these interactions. VM and DL conducted mass spectrometry (MS) for the CARM1 and ALIX interactomes and the determination of arginine residues that are methylated in ALIX. SH, SS, YY, RDa, and SR were responsible for cloning. YY, CV, and LA assessed the specificity of ALIX interaction with PRMTs. SH and SS biochemically characterized the CARM1-ALIX interaction. SH, SS, CC, RS, AEM, VM, and DL studied ALIX methylation. JV and NR performed experiments and analysis to identify the methyl-sensitive ALIX partner, with MS analysis done by LB and YC; SH validated these partners by western blot. RG performed the AlphaFold2 predictions and their analyses. SH analyzed the effects of ALIX mutation on cytokinesis. CC, RS, AEM, AP, RDi, JT and AE provided reagents and expertise. SH, SS, AE, and TD designed the study; AP, RDi, JT and AE being also involved in designing the studies regarding cytokinesis. SH, AE and TD wrote the manuscript and created the figures and tables. All authors reviewed and edited the manuscript. TD provided overall supervision and secured funding.

## Competing interests

The authors declare no competing interests.

## Notes

### Competing Interest Statement

The authors have declared no competing interest.

