## Supplementary Figures 1-9 for "CARM1-mediated methylation controls interactions of ALIX with key partners important for cytokinesis"

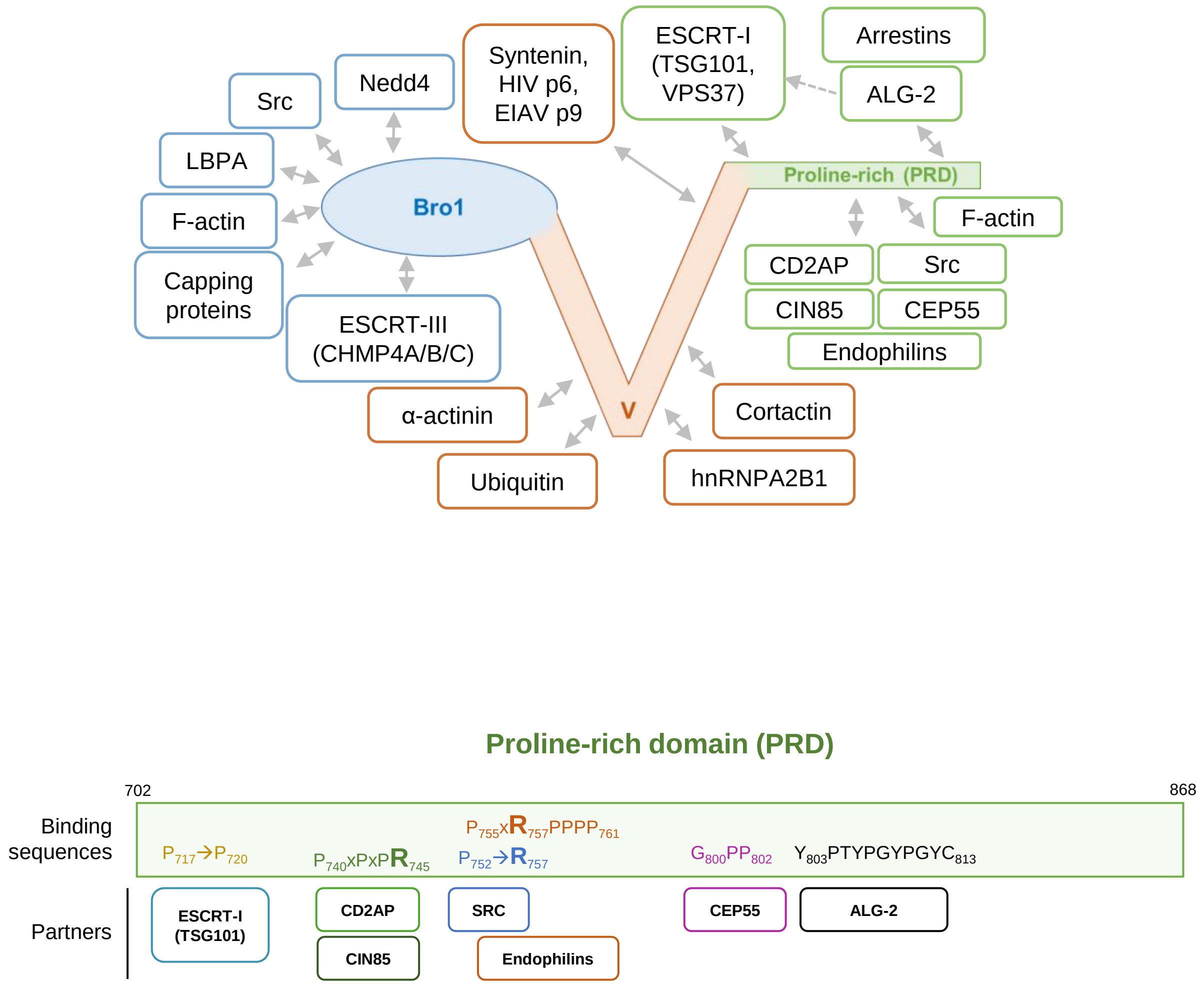

**Supplementary Figure 1 | ALIX domains and partners.** Schematic representation of ALIX Bro1 domain (shown in blue), V domain (shown in orange) and PRD (shown in green) with their partners (top panel). Zoom on ALIX PRD with its partners and binding sequences (bottom panel).

**HCC1187**
0.1% NP-40

| protein name | MW (kDa) | IgG M | IP Ab1 CARM1 |
|---|---|---|---|
| CARM1 | 65.9 | 4 | 67 |
| ALIX | 96 | 0 | 48 |
| QARS1 | 87.8 | 5 | 16 |
| RBBP8 | 101.9 | 0 | 10 |
| PCMT1 | 24.6 | 0 | 7 |

| protein name | MW (kDa) | IgG R | IP Ab2 CARM1 |
|---|---|---|---|
| IQGAP1 | 189.3 | 0 | 106 |
| CARM1 | 65.9 | 0 | 68 |
| ALIX | 96 | 0 | 54 |
| ERC1 | 128 | 0 | 36 |
| AMOT | 118 | 0 | 31 |

| protein name | MW (kDa) | IgG R | IP Ab3 CARM1 |
|---|---|---|---|
| CARM1 | 65.9 | 0 | 61 |
| ALIX | 96 | 0 | 48 |
| GAB2 | 74.5 | 0 | 31 |
| SPAG9 | 146.2 | 0 | 11 |
| ANKRD10 | 44.8 | 0 | 11 |

**HCC1187**
**1% NP-40**

| protein name | MW (kDa) | IgG M | IP Ab1 CARM1 |
|---|---|---|---|
| MYH10 | 229.0 | 5 | 75 |
| CARM1 | 65.9 | 11 | 72 |
| ALIX | 96 | 0 | 48 |
| RBBP8 | 101.9 | 0 | 28 |
| SVIL | 247.7 | 0 | 19 |

| protein name | MW (kDa) | IgG R | IP Ab2 CARM1 |
|---|---|---|---|
| IQGAP1 | 189.3 | 0 | 70 |
| CARM1 | 65.9 | 6 | 67 |
| ALIX | 96 | 0 | 46 |
| ERC1 | 128.1 | 0 | 40 |
| KCTD20 | 47.5 | 0 | 33 |

| protein name | MW (kDa) | IgG R | IP Ab3 CARM1 |
|---|---|---|---|
| CARM1 | 65.9 | 11 | 63 |
| ALIX | 96 | 0 | 43 |
| GAB2 | 74.5 | 0 | 29 |
| TRIM21 | 54.2 | 2 | 21 |
| SPAG9 | 146.2 | 0 | 8 |

**BT-549**
0.1% NP-40

| protein name | MW (kDa) | IgG M | IP Ab1 CARM1 |
|---|---|---|---|
| CARM1 | 65.9 | 0 | 120 |
| ALIX | 96 | 0 | 104 |
| P4HB | 57.1 | 0 | 22 |
| RBBP8 | 101.9 | 0 | 11 |
| PABPC1 | 70.7 | 9 | 20 |

| protein name | MW (kDa) | IgG R | IP Ab2 CARM1 |
|---|---|---|---|
| ERC1 | 128.1 | 3 | 135 |
| ALIX | 96 | 0 | 125 |
| CARM1 | 65.9 | 5 | 120 |
| IQGAP1 | 189.3 | 17 | 125 |
| KIDINS220 | 196.5 | 0 | 67 |

| protein name | MW (kDa) | IgG R | IP Ab3 CARM1 |
|---|---|---|---|
| ALIX | 96 | 0 | 107 |
| CARM1 | 65.9 | 5 | 85 |
| SPAG9 | 146.2 | 0 | 43 |
| ABCD3 | 75.5 | 0 | 41 |
| ANXA1 | 38.7 | 7 | 27 |

**HeLa 0.1%**
NP-40

| protein name | MW (kDa) | IgG M | IP Ab1 CARM1 |
|---|---|---|---|
| CARM1 | 65.9 | 5 | 101 |
| ALIX | 96 | 3 | 76 |
| CASP14 | 27.7 | 0 | 14 |
| RBBP8 | 101.9 | 0 | 12 |
| TGM3 | 76.6 | 0 | 8 |

| protein name | MW (kDa) | IgG R | IP Ab2 CARM1 |
|---|---|---|---|
| IQGAP1 | 189.3 | 27 | 203 |
| ERC1 | 128.1 | 6 | 173 |
| CARM1 | 65.9 | 11 | 119 |
| ALIX | 96 | 8 | 98 |
| KIDINS220 | 196.5 | 0 | 71 |

| protein name | MW (kDa) | IgG R | IP Ab3 CARM1 |
|---|---|---|---|
| ALIX | 96 | 8 | 86 |
| CARM1 | 65.9 | 11 | 80 |
| BIRC6 | 560.3 | 0 | 51 |
| SAMHD1 | 72.2 | 0 | 51 |
| CEP128 | 128 | 0 | 41 |

**Supplementary Figure 2 | ALIX is a main partner of CARM1.** Peptide counts detected by mass-spectrometry for the top five retrieved proteins in CARM1 immunoprecipitates (IP) using three different CARM1 antibodies (Ab1, left tables; Ab2, middle tables; Ab3, left tables) in various cell lines. IgG were used as a control. Molecular weight (MW) are indicated. Experiments were performed in 0.1% NP-40 except one with 1% NP-40 as indicated. The peptide counts for CARM1 and ALIX (0.1% NP-40) are those reported in Fig. 1a.

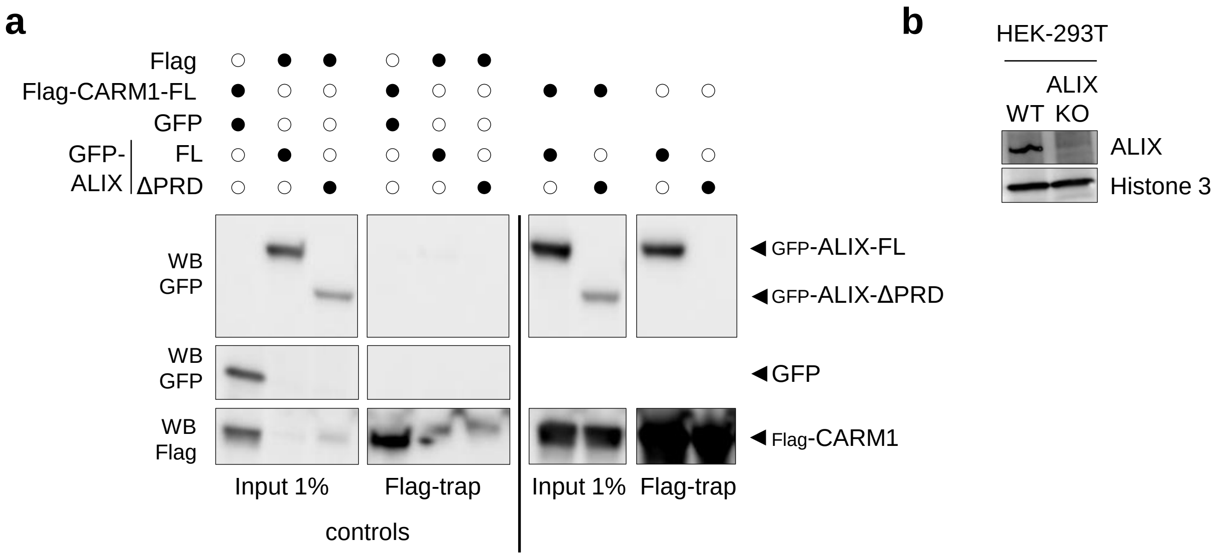

**Supplementary Figure 3 | ALIX PRD is required for CARM1 binding. a** HEK293 ALIX KO cells were transfected with the constructs as indicated. Flag-CARM1 was immunoprecipitated using a Flag-trap and its interaction with the different GFP-ALIX proteins was assessed by western blot (images are representative of three independent experiments). Blots containing the negative controls (left panels) or the interaction between Flag-CARM1 and the two GFP-ALIX proteins (right panels) are separated due to the presence of additional lanes not relevant to this study. **b** Validation of ALIX KO by western blot in HEK-293T ALIX KO cells. Histone 3 antibody was used as a loading control.

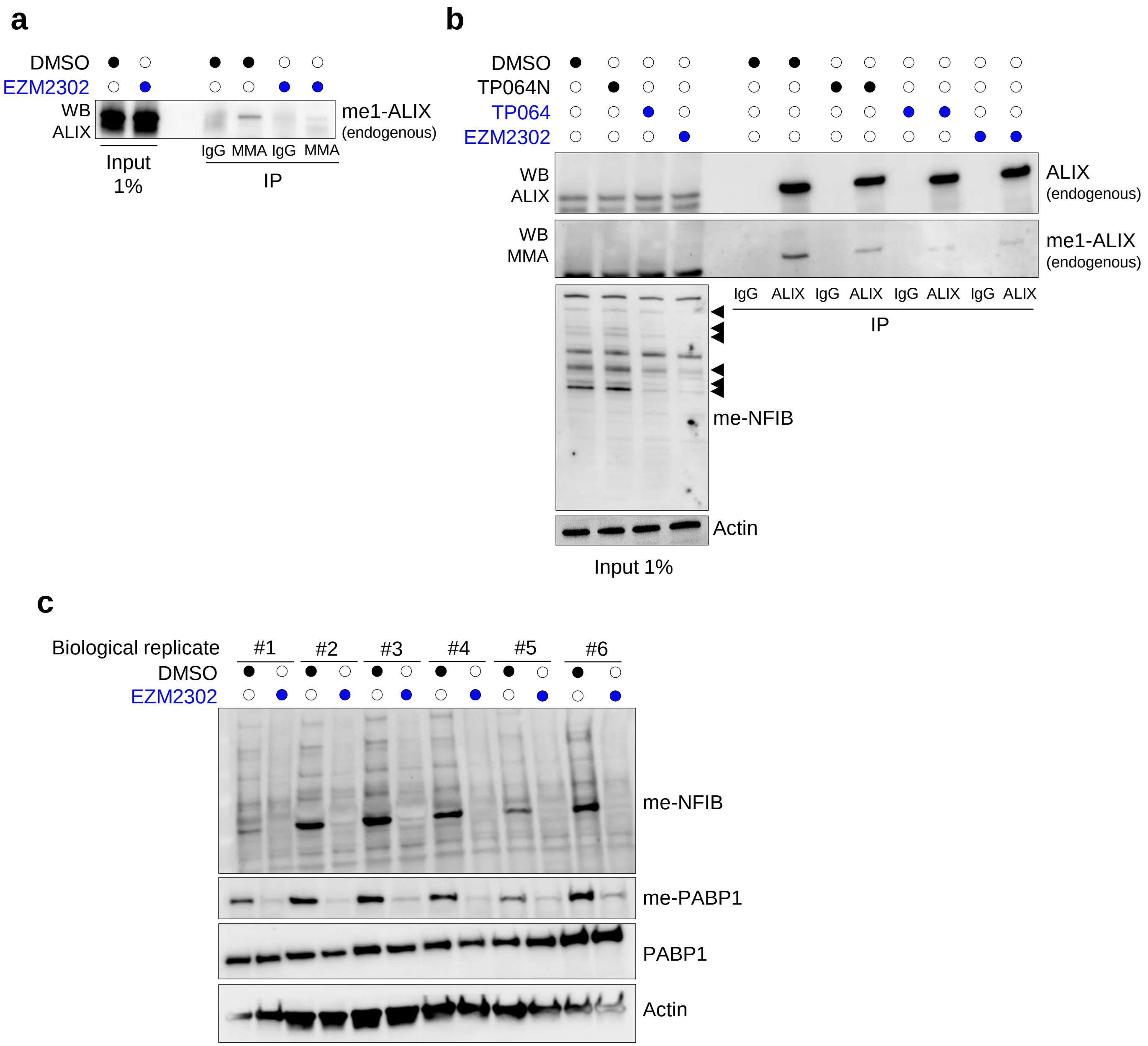

**Supplementary Figure 4 | CARM1 methylates ALIX in cells. a** HeLa cells treated with 0.5 µM EZM2302 or with DMSO for 48 h before immunoprecipitation (IP) using monomethylarginine (MMA) or IgG antibodies. ALIX monomethylation was assessed by western blot (*N* = *3*). Similar results were obtained in BT-549 cells (data not shown). Actin was used as a loading control. **b** HeLa cells treated with 1 µM CARM1 inhibitors (TP064, EZM2302), 1 µM control compound (TP064N) or with DMSO for 48 h were subjected to IP using ALIX or IgG antibodies. ALIX monomethylation was assessed by western blot using anti-monomethylarginine (MMA) antibody (upper panel, *N* = *2*). In the lower panels, the efficacy of CARM1 inhibition was confirmed by western blotting with anti-meNFIB antibody[61]. **c** Western blot confirming CARM1 inhibition using anti-meNFIB and anti-mePABP1 antibodies in the protein lysates used in Fig. 3d. The anti-meNFIB antibody detects methylated NFIB, as well as other methylated proteins[61], and was used here as a pan-ADMA antibody. Arrows indicate a decrease of me2a proteins **(b)**.

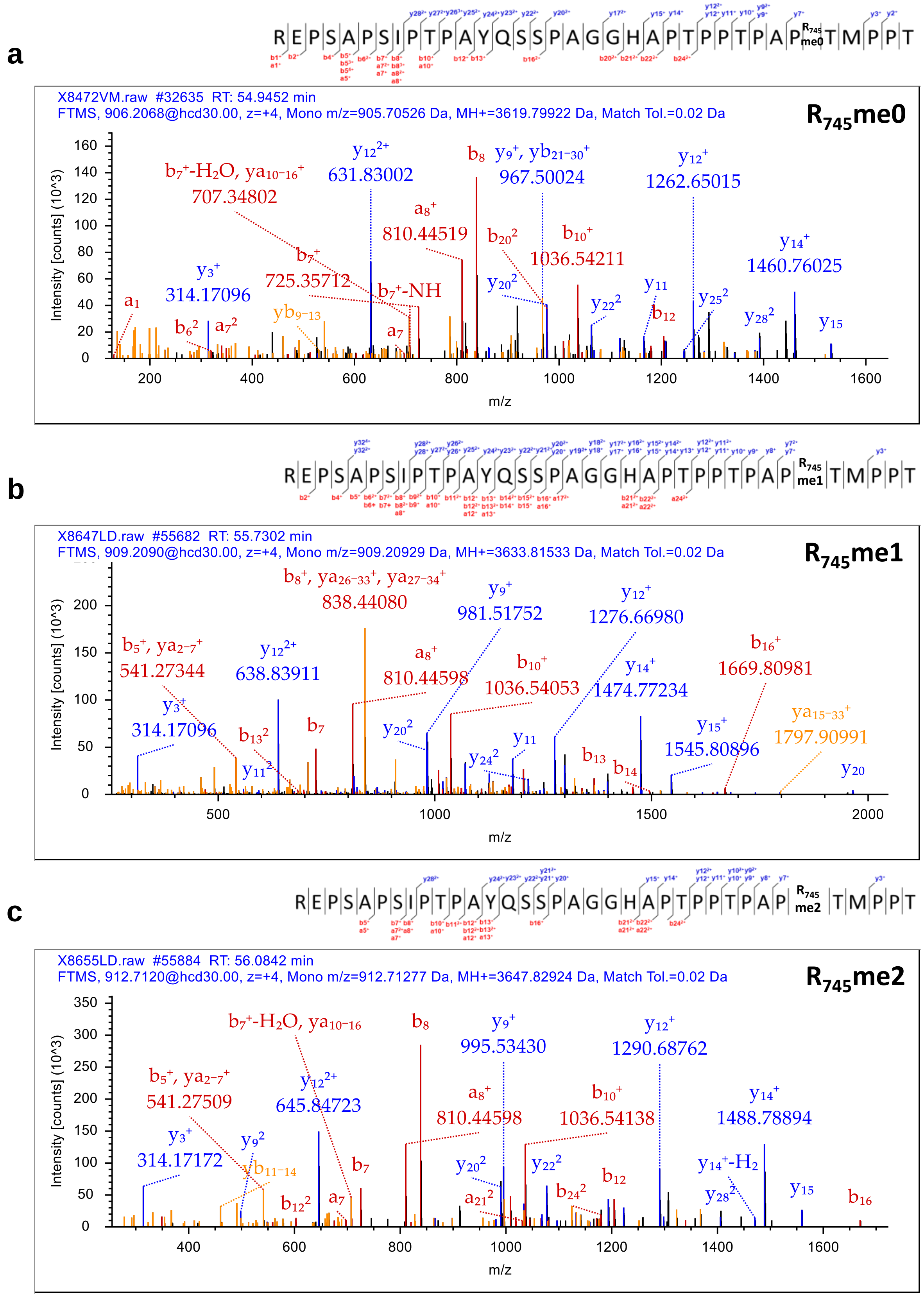

**Supplementary Figure 5 | Representative MS/MS spectra of unmodified, methylated and dimethylated ALIX $R_{745}$. a-c** Fragmentation spectra derived from Lysarginase digested ALIX. The unmodified (me0) **(a)**, monomethylated (me1) **(b)** and dimethylated (me2) **(c)** peptide sequences and observed ions are indicated on top of spectra. Singly, doubly and triply charges a, b and y ions, as well as ions corresponding to neutral losses of water ($H_2O$) and $NH_3$ groups, M, parent ion and internal fragments are shown. Spectrums are representative of four independent experiments.

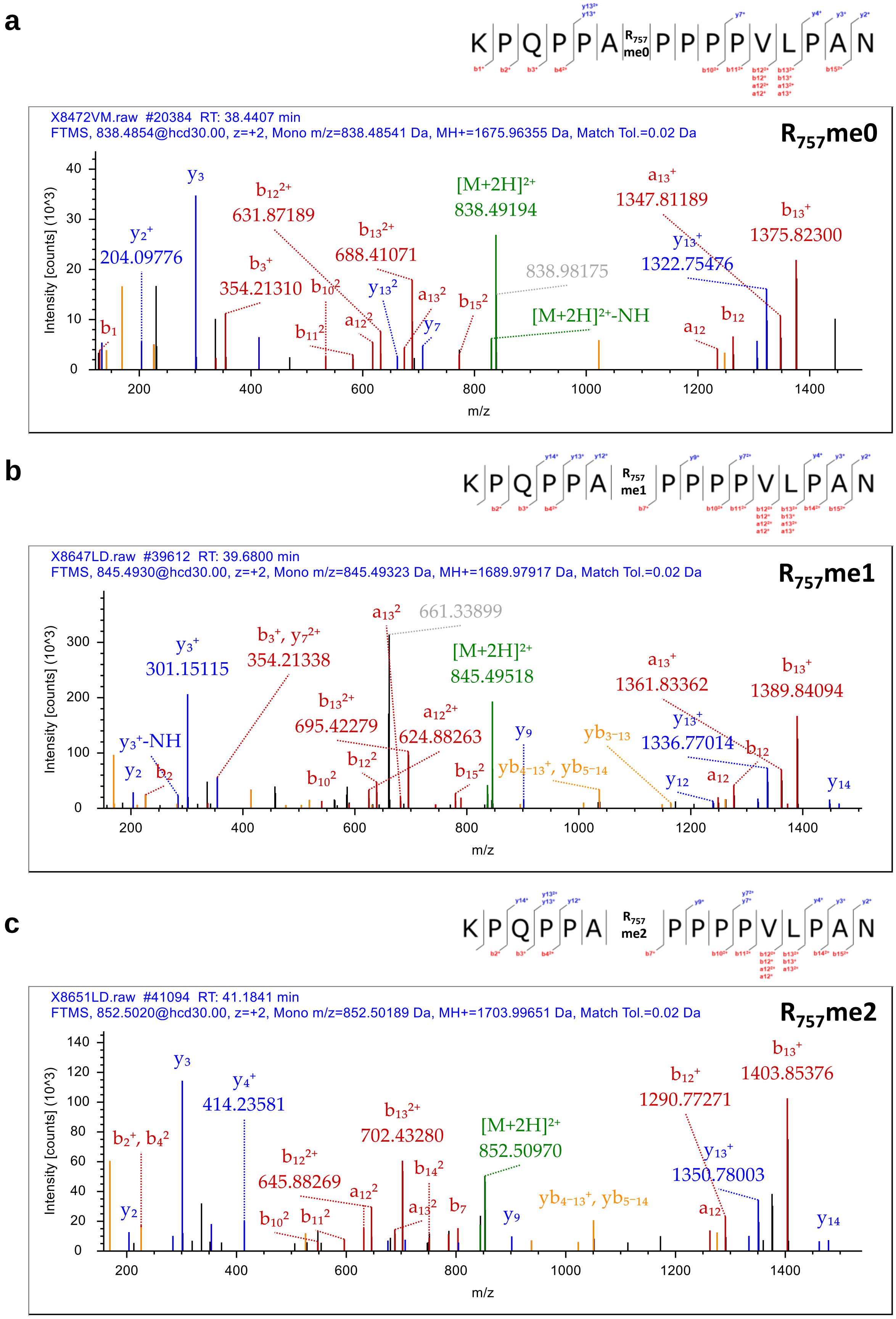

**Supplementary Figure 6 | Representative MS/MS spectra of unmodified, methylated and dimethylated ALIX $R_{757}$. a-c** Fragmentation spectra derived from Lysarginase digested ALIX. The unmodified (me0) **(a)**, monomethylated (me1) **(b)** and dimethylated (me2) **(c)** peptide sequences and observed ions are indicated on top of spectra. Singly, doubly and triply charges a, b and y ions, as well as ions corresponding to neutral losses of water ($H_2O$) and $NH_3$ groups, M, parent ion and internal fragments are shown. Spectrums are representative of five independent experiments.

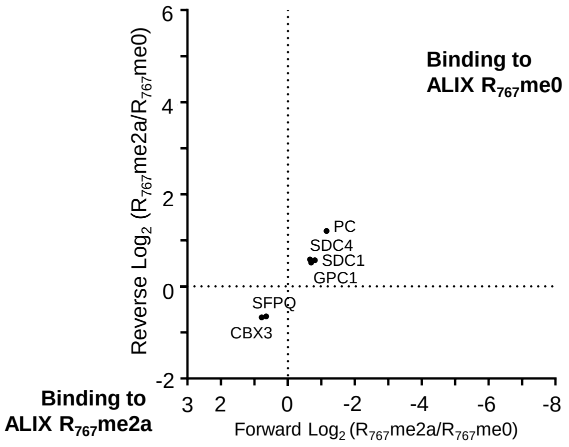

**Supplementary Figure 7 | Peptide pull down to identify proteins interacting specifically to unmethylated or asymmetrically dimethylated $R_{767}$ of ALIX.** Peptide pull down assays were performed with protein lysates from MDA-MB-231 cells. ALIX binding partners were identified using SILAC-based quantitative proteomics screen. A scatter plot of the $Log_2$ transformed protein-normalized methyl-sensitive peptide SILAC ratios. The x-axis shows the $Log_2$ ratio of proteins binding to asymmetrically dimethylated (me2a) versus unmethylated (me0) arginine residues (Forward). The y-axis shows the $Log_2$ ratio of a label-swap replicate experiment (Reverse). Specific interactors of unmethylated and asymmetric dimethylated ALIX reside in the upper right and in the lower left quadrant, respectively. The experiment was done twice with similar results.

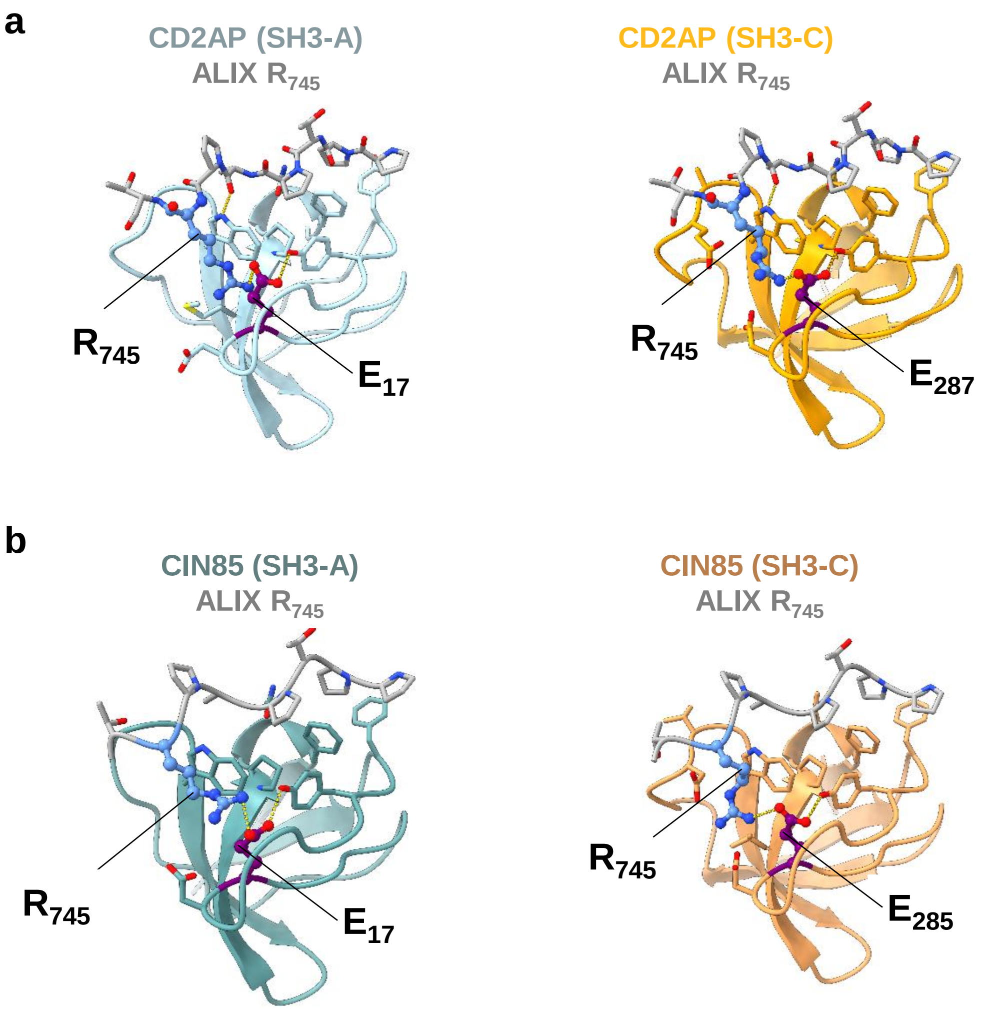

**Supplementary Figure 8 | The arginine residue $R_{745}$ within the PxPxPR motif of ALIX forms salt bridges with glutamic acid residues in SH3 domains of CD2AP and CIN85.** **a, b** AlphaFold2 based structural models of the complexes between $R_{745}$ of ALIX peptides and SH3-A (left) and SH3-C (right) domains of CD2AP **(a)** or CIN85 **(b)**. $R_{745}$ forms key salt bridges with conserved glutamic acid in CD2AP SH3-A ($E_{17}$), CD2AP SH3-C ($E_{287}$), CIN85 SH3-A ($E_{17}$), CIN85 SH3-C ($E_{285}$). Details on peptide lengths, SH3 domains boundaries, and model confidence scores are provided in Supplementary Fig. 9.

| SH3 domains residues | ALIX Peptide residues | pLDDT | pTM score | ipTM score | combinedTM score |
|---|---|---|---|---|---|
| **CD2AP (SH3-A)** 1-55 | $R_{745}$ 729-750 | **84.30** | **0.72** | **0.52** | **0.56** |
| **CD2AP (SH3-B)** 110-166 | $R_{745}$ 729-750 | **86.35** | **0.74** | **0.58** | **0.61** |
| CD2AP (SH3-C) 270-330 | $R_{745}$ 729-750 | 84.46 | 0.73 | 0.54 | 0.58 |
| CIN85 (SH3-A) 1-58 | $R_{745}$ 738-751 | 89.62 | 0.78 | 0.57 | 0.61 |
| **CIN85 (SH3-B)** 101-157 | $R_{745}$ 738-751 | **92.46** | **0.81** | **0.68** | **0.71** |
| **CIN85 (SH3-C)** 270-336 | $R_{745}$ 738-751 | **89.21** | **0.78** | **0.68** | **0.70** |
| **Endophilin A2 (SH3)** 300-368 | $R_{757}$ 746-770 | **82.5** | **0.72** | **0.51** | **0.55** |

**Supplementary Figure 9 | Confidence scores associated with the AlphaFold2 derived models.**
