## Supplementary Material Tables 1-4 for "CARM1-mediated methylation controls interactions of ALIX with key partners important for cytokinesis"

### Table 1: List of plasmids and primers

| Primers | Used to generate the construct | Target sequence (5' → 3') | Application |
|---|---|---|---|
| CARM1-Δ15-deletion-F | Flag-CARM1-Δ15 | CCACAACAACCTGATTCCTTTA GGGTCCTCCG | Mutagenesis |
| CARM1- Δ15-deletion-R | | CGGAGGACCCTAAAGGAATCA GGTTGTTGTGG | Mutagenesis |
| Flag-CARM1-HindIII-F | Flag-CARM1-PH*, Flag-CARM1-ΔCter* | CTACGAAGCTTATGGACTACAA AGACGATGACG | Cloning |
| Flag-CARM1-PH-EcoRI-R | Flag-CARM1-PH* | GCTACGGAATTCCTAGTGGCCC CGGCAGGTTTTC | Cloning |
| Flag-CARM1-catalytic-HindIII-F | Flag-CARM1-Catalytic* | CTACGAAGCTTATGGACTACAA AGACGATGACGACAAGCACAC CCTGGAGCGGTCTG | Cloning |
| Flag-CARM1-PH-catalytic-EcoRI-R | Flag-CARM1-ΔCter*, Flag-CARM1-Catalytic* | GCTACGGAATTCCTAGCTGCTG AGGTTGTAGGTGC | Cloning |
| Flag-CARM1-catalytic-Cterm-HindIII-F | Flag-CARM1-ΔPH* | CTACGAAGCTTATGGACTACAA AGACGATGACGACAAGCACAC CCTGGAGCGGTCTG | Cloning |
| Flag-CARM1-catalytic-Cterm-EcoRI-R | | GCTACGGAATTCCTAGCTCCCG TAGTGCATGGT | Cloning |
| GST-BamHI-F | PfastBac-GST | TAGTCAGGATCCATGTCCCCTAT ACTAGGTTATTG | Cloning |
| GST-EcoRI-R | | TGACTAGAATTCCGATCCACGC GGAACCAGATCCG | Cloning |
| CARM1-EcoRI-F | pFastBac-GST-CARM1-FL | TGCTAGGAATTCATGGACAAG GCAGCGGCGGCGG | Cloning |
| CARM1-XbaI-R | | CTAGCATCTAGACTAGCTCCCG TAGTG CATGG | Cloning |
| Flag-PRMT1-F | Flag-PRMT1 | CATGGACTACAAAGACGATGA CGACAAGATGGCGGCAGCCG AGGCC | Cloning |
| Flag-PRMT1-R | | GATCAGCGGGTTTAAACGGGC CCTCTAGATCAGCGCATCCG GTAGTCGG | Cloning |
| Flag-HiFi-F | Flag-PRMT1 | CTAGAGGGCCCGTTTAAACC | Cloning – HiFi assembly |
| Flag-HiFi-R | Flag-PRMT1 | CTTGTCGTCATCGTCTTTG | Cloning – HiFi assembly |

| | | | |
|---|---|---|---|
| Flag-PRMT2-F | Flag-PRMT2 | GCAACATCAGGTGACTGTC | Cloning |
| Flag-PRMT2-R | | TCTCCAGATGGGGAAGAC | Cloning |
| mcherry-ALIX-V-stopcodon-F | mcherry-ALIX-V | CGGAAGACAGAAAGAGATTAACTCTTAAAGGAC | Mutagenesis |
| mcherry-ALIX-V-stopcodon-R | | GTCCTTTAAGAGTTAATCTCTTTCTGTCTTCCG | Mutagenesis |
| mcherry-ALIX-PRD-EcoRI -F | mcherry-ALIX-PRD | TGCTAGGAATTCAGATGAACTCTTAAAGGACTTGCAAC | Cloning |
| mcherry-ALIX-PRD-BamHI-R | | CTAGCAGGATCCCTACTGCTGTGGATAGTAAGACTGC | Cloning |
| GFP-ALIX-BamHI-F | GFP-ALIX-ΔPRD | CTACGGGATCCATGGTGAGCAAGGGCGAGGAGCTG | Cloning |
| GFP-ALIX_1-702-XmaI-R | | GCTACGCCCGGGCTATCTTTCTGTCTTCCGTGC | Cloning |
| GFP-ALIX-FL-$R_{745}K$-F | GFP-ALIX-3K | CCAACTCCAGCGCCAAAAACCATGCCGCCTAC | Mutagenesis |
| GFP-ALIX-FL-$R_{745}K$-R | | GTAGGCGGCATGGTTTTTGGCGCTGGAGTTGG | Mutagenesis |
| GFP-ALIX-FL-$R_{757}K$-F | | CCCAGCCCCCAGCCAAGCCTCCACCACCTG | Mutagenesis |
| GFP-ALIX-FL-$R_{757}K$-R | | CAGGTGGTGGAGGCTTGGCTGGGGGCTGGG | Mutagenesis |
| GFP-ALIX-FL-$R_{767}K$-F | | CTGTGCTTCCAGCAAATAAAGCTCCTTCTGCTACTGCTCC | Mutagenesis |
| GFP-ALIX-FL-$R_{767}K$-R | | GGAGCAGTAGCAGAAGGAGCTTTATTTGCTGGAAGCACAG | Mutagenesis |
| GB1-His-ALIX-$PRD_{703-800}$ _$R_{745}K$-F | ALIX-$PRD^{703-800}$-2K, ALIX-$PRD^{703-800}$-3K | CCAACGCCGGCCCCAAAGACGATGCCGCCAACC | Mutagenesis |
| GB1-His-ALIX-$PRD_{703-800}$ _$R_{745}K$-R | | GGTTGGCGGCATCGTCTTTGGGGCCGGCGTTGG | Mutagenesis |
| GB1-His-ALIX-$PRD_{703-800}$ _$R_{757}K$-F | | CCGCAACCACCAGCGAAGCCGCCACCACCAGTTC | Mutagenesis |
| GB1-His-ALIX-$PRD^{703-800}$ _$R_{757}K$-R | | GAACTGGTGGTGGCGGCTTCGCTGGTGGTTGCGG | Mutagenesis |
| GB1-His-ALIX-$PRD_{703-800}$ _$R_{767}K$-F | | GTTCTCCCAGCCAATAAGGCCCCAAGTGCGACC | Mutagenesis |
| GB1-His-ALIX-$PRD_{703-800}$ _$R_{767}K$-R | | GGTCGCACTTGGGGCCTTATTGGCTGGGAGAAC | Mutagenesis |
| PB5-ALIX | iON-CAG-GFP-ALIX-WT, iON-CAG-GFP-ALIX-3K | CAACTAGAAGGCACTAAAGTCGGATCCTCGCTACTGCTGTGGATAGTAAGACTG | Cloning |

| | | | |
|---|---|---|---|
| PB5-GFP | iON-CAG-GFP | CAACTAGAAGGCACTAAAGTCGGATCCTCGTTACTTGTACAGCTCGTCCATGC | Cloning |
| PB3-GFP | iON-CAG-GFP-ALIX-WT, iON-CAG-GFP-ALIX-3K, iON-CAG-GFP | GGGTTAATTAAGTGATTAGCCCGGGCCTAGGCCACCATGGTGAGCAAGGGCGAGGAGCTG | Cloning |

F, forward primer, R, reverse primer. *all constructs marked with an Asterix were produced using the pcDNA3.1(+)-Flag-CARM1-FL plasmid.

**Table 2: List of siRNA**

| siRNA | Target Sequence 5'→3' | Company | Reference |
|---|---|---|---|
| AllStars Negative Control siRNA | - | Qiagen | 0001027281 |
| ALIX (for siRNA-resistant plasmids) | CCTGGATAATGATGAAGGA | Synthesized by Eurofins | Ref[26] |

**Table 3: List of antibodies used**

| Target | Company | Reference | Host species | Application | Dilution |
|---|---|---|---|---|---|
| ALIX | Cell Signaling Technologies | 92880 | R | WB | 1/1,000 |
| ALIX | Bethyl | A302-938A | R | IP | 2 µg/mg protein lysate |
| CAPZA1 | Proteintech | 11806-1-AP | R | WB | 1/1,000 |
| CAPZB | Proteintech | 25043-1-AP | R | WB | 1/1,000 |
| CARM1 (Ab2) | Cell Signaling Technologies | 3379 | R | WB, IP | 1/1,000, 2 µg/mg protein lysate |
| CARM1 (Ab1) | Cell Signaling Technologies | 12495 | M | WB, IP | 1/1,000, 2 µg/mg protein lysate |
| CARM1 (Ab 3) | Bethyl | A300-421A | R | IP | 2 µg/mg protein lysate |
| CD2AP | Proteintech | 51046-1-AP | R | WB | 1/1,000 |
| Endophilin-A2 | ThermoScientific (kind gift from L. Johannes, I. Curie) | PA-5-22292 | R | WB | 1/1,000 |
| Flag (clone M2) | Sigma-Aldrich | F3165 | M | WB, IP | 1/2,000, 2µg/mg lysate |
| GAPDH | Cell Signaling Technologies | 2118 | R | WB | 1/1,000 |
| GFP | CurieCore Antibody platform, I. Curie | A-P-R#006 | R | WB, IP | 1/1000, 2 µg/mg protein lysate |
| Histone H3 | Cell signaling | 9717 | R | WB | 1/1,000 |
| mcherry | Antibody platform – I. Curie | A-P-R#13 | R | WB, IP | 1/2,500, 2 µg/mg protein lysate |
| meNFIB | Homemade, kind gift from Dr M. Bedford (MD Anderson, USA) (ref[61]) | - | R | WB | 1/1,000 |
| Methyl-PABP1 (asymmetric R455/R460) | Cell Signaling Technologies | 3505 | R | WB | 1/1,000 |
| Mono-methyl arginine | Cell Signaling Technologies | 8015 | R | WB, IP | 1/1,000, 2 µg/mg protein lysate |

| | | | | | |
|---|---|---|---|---|---|
| PABP1 | Cell Signaling Technologies | 4992 | R | WB | 1/1,000 |
| Anti-Mouse IgG-HRP | Interchim | 115-035-062 | G | WB | 1/20,000 |
| Anti-Mouse light chain HRP | Upstate | AP200P | G | WB | 1/20,000 |

R, rabbit; M, mouse; H, human; H, goat

**Table 4: List of synthetic peptides**

| Peptide name | Sequence | Molecular weight (Da) | Application |
|---|---|---|---|
| Biotin-ALIX-$R_{745}$ | Biotin-KK-$T_{738}$PPTPAP**$R_{745}$**TMPP$T_{750}$-KKK-$NH_2$ | 2229 | Pulldown |
| Biotin-ALIX-$R_{745}$-ADMA | Biotin-KK-$T_{738}$PPTPAP(**$R_{745}$ADMA**)TMPP$T_{750}$-KKK-$NH_2$ | 2257 | Pulldown |
| Biotin-ALIX-$R_{757}$ | Biotin-KKK-$P_{752}$QPPA**$R_{757}$**PPPPVLP$A_{765}$-KK-$NH_2$ | 2300 | Pulldown |
| Biotin-ALIX-$R_{757}$-ADMA | Biotin-KKK-$P_{752}$QPPA(**$R_{757}$ADMA**)PPPPVLP$A_{765}$-KK-$NH_2$ | 2327 | Pulldown |
| Biotin-ALIX-$R_{767}$ | Biotin-KKK-$P_{761}$VLPAN**$R_{767}$**APSATAP$S_{775}$-KKK-$NH_2$ | 2443 | Pulldown |
| Biotin-ALIX-$R_{767}$-ADMA | Biotin-KKK-$P_{761}$VLPAN(**$R_{767}$ADMA**)APSATAP$S_{775}$-KKK-$NH_2$ | 2470 | Pulldown |
